# Dietary fructose promotes anxiety through impairment of microglia physiology

**DOI:** 10.64898/2026.10.23.761346

**Authors:** Adeline Coursan-Crave, Thomas Devos, Mathilde Huart, Sandrine Ellero-Simatos, Maguelonne Peffault de Latour, Shingo Nakajima, Stéphanie Fulton, Sophie Layé, Thierry Alquier, Charlotte Madore, Véronique Douard, Xavier Fioramonti

**Affiliations:** Univ. Bordeaux, INRAE, Bordeaux INP, NutriNeuro, UMR 1286, F-33000 Bordeaux, France; Toxalim (Research Center in Food Toxicology), INRAE UMR 1331, ENVT, INP-Purpan, Université de Toulouse, 31300, Toulouse; Centre de Recherche du Centre Hospitalier de l’Université de Montréal (CRCHUM), Montreal, Quebec, Canada; Université Paris-Saclay, INRAE, AgroParisTech, MICALIS Institute, Jouy-en-Josas, France

**Keywords:** Fructose, Glut5, microglia, anxiety.

## Abstract

Excessive dietary fructose has been linked to metabolic and mood disorders but the underlying brain mechanisms remain unclear. In this study, we tested whether microglia, the brain’s resident immune cells, mediate fructose-induced anxiety. Adult C57BL/6J mice and mice knock-down for the fructose transporter Glut5 selectively in microglia (Slc2a5flox/flox:Tmem119-Cre-positive mice) were exposed to 25% fructose in drinking water for 8 weeks. We assessed metabolic parameters, anxiety- and depression-like behaviors using a battery of behavioral tests. We then performed RNA sequencing in several brain regions, including the hippocampus to characterize fructose-induced transcriptional changes. Microglial responsiveness to lipopolysaccharide (LPS) treatment was assessed *in vitro* and *in vivo* using immunohistochemistry (microglia morphology) and RT-qPCR (cytokine gene expression). Fructose feeding induced marked metabolic alterations and robust anxiety-like behavior in male mice, together with memory deficits, whereas females showed milder metabolic changes and memory impairment without clear emotional disturbances. RNA sequencing revealed that fructose alters microglial gene-expression profiles in the hippocampus. Microglial Glut5 knockdown prevented fructose-induced anxiety-like behavior in the novelty-suppressed feeding test, indicating that microglial fructose uptake is required for this effect. At the molecular level, physiological fructose exposure did not markedly increase microglial cytokine expression but blunted the cytokine response to lipopolysaccharide, consistent with a hypo-reactive microglial state. Overall, these findings identify microglia as a critical mediator of fructose-induced anxiety and suggest that chronic fructose exposure promotes a maladaptive microglial phenotype that disrupts brain circuit function, with potential implications for diet-related mental health risk.

**HIGHLIGHTS:**

- Fructose feeding induces anxiety in adult male mice but not in females
- Fructose feeding alters hippocampal gene expression representative of microglial signature
- Effect of fructose feeding on anxiety depend on microglial fructose uptake
- Fructose directly blunts the microglial response to LPS

## BACKGROUND

The impact of unbalanced nutrition on brain function has become a major public health concern. Among dietary factors, excessive fructose consumption has been highlighted by international health agencies, including WHO and ANSES, as potentially deleterious (Tappy et al., 2018). Since the 1960s, fructose has been widely incorporated into processed foods and beverages, leading to a marked increase in intake—from less than 5 g/day historically to approximately 50–80 g/day in developed countries(Tappy and Lê, 2010). In Europe, the abolition of sugar production quotas in 2017 may further contribute to this rise.

Most studies have focused on the deleterious effects of fructose on metabolic disorders, including obesity and type 2 diabetes (T2D). More recently, both human and animal studies suggest that excessive fructose intake is also associated with impairments in brain function, including anxiety, depressive-like symptoms, and cognitive deficits (Spagnuolo et al., 2020; Gillespie et al., 2023; Barma et al., 2026). However, the cellular and molecular mechanisms underlying these neurobehavioral effects remain poorly understood. While altered glucose homeostasis associated with T2D has been linked to mood disorders (Martin et al., 2021; Fulton et al., 2022), it remains unclear whether fructose exerts direct effects on the brain independent of systemic metabolic alterations.

Emerging evidence indicates that fructose can access the brain and may be locally metabolized (Hwang et al., 2015; Wang et al., 2025). Notably, the fructose transporter GLUT5 is predominantly expressed in microglia within the central nervous system (Zhang et al., 2014). Microglia play a central role in neuroimmune regulation, modulating neuroinflammation through cytokine release and shaping neural circuits via synaptic pruning and phagocytosis (Colonna and Butovsky, 2017). Recent studies suggest that fructose may influence microglial function (Ting, 2024a; Wang et al., 2025), but direct evidence linking fructose exposure to microglial alterations driving behavioral outcomes is still lacking. In particular, recent studies have suggested that early-life high fructose exposure impairs microglial phagocytosis and disrupts neurodevelopment in a GLUT5-dependent manner, leading to anxiety-like behaviors in adolescence (Wang et al., 2025). However, the consequences of chronic fructose intake during adulthood on microglial function and the regulation of emotional behavior remain largely unexplored despite the fact that excessive fructose consumption is also prevalent in adult populations. To address this gap, we combined complementary *in vivo* and *in vitro* approaches to investigate the impact of fructose on microglial function and its contribution to the development of emotional disorders.

## MATERIALS AND METHODS

### Animals

Mice (8-12 weeks old) were group-housed in conventional housing cages at the NutriNeuro animal facility (CIRCE Bordeaux) and maintained on a 12-hour light-dark cycle with ad libitum access to food and water. Fructose fed animals received fructose 25% in their drinking water for 8 weeks. In a subgroup of experiments, animals were injected intraperitoneally with lipopolysaccharide (LPS; 250 µg/kg) 3 or 24 hours before euthanasia to perform RT-qPCR on isolated microglia or immunohistochemistry, respectively. All experimental procedures were conducted in accordance with the European directive 2010/63/UE and approved by the French Ministry of Research and local ethics committees (APAFIS# 13272 and APAFIS# 19870).

### Construction of Glut5-KD mice

Cryopreserved sperm for the C57BL/6N-Slc2a5tm1a(EUCOMM)Wtsi/Ieg (ID: 07602) strain were obtained from the European Mutant Mouse Archive. Sperm carrying the tm1a-KO first/pre-conditional allele and oocytes from wild-type (WT) donor C57BL/6N females were used to perform in vitro fertilization at the CRYME core facility of Bordeaux Neurocampus. Heterozygous mice were then crossed to the FLPo delete strain B6.129S4-Gt(ROSA)26Sortm2(FLP*)Sor/N (JAX stock number 012930) to eliminate the lacZ reporter and neo cassette and generate a clean conditional floxed allele tm1c (Slc2a5flox/flox). Homozygous Slc2a5flox/flox mice with loxP sites flanking exon 6 of the gene were crossed to C57BL/6-Tmem119^em1(cre/ERT2)Gfng^/N to generate Slc2a5flox/flox cre negative or Slc2a5flox/flox:Tmem119-Cre-positive mice (always heterozygous for Tmem119-cre). 5-weeks old Slc2a5flox/flox cre negative and Slc2a5flox/flox:Tmem119-Cre-positive mice were treated daily with tamoxifen (75 mg/kg diluted in corn oil, IP) for 5 consecutive days to generate WT (Slc2a5flox/flox cre negative) or Glut5-KD (Slc2a5flox/flox cre negative and Slc2a5flox/flox:Tmem119-Cre-positive) mice which were used 3 weeks post-tamoxifen treatment for experiments.

### Behavioral tests

Behavioral tests were performed after 8 weeks of diet while animals were maintained under the same diet during the tests. Animals were tested in different anxiety-, depressive-like as well as memory behavioral paradigms as previously described (Zemdegs et al., 2016, 2019; Martin et al., 2024). Briefly, the elevated plus maze (EPM) test was performed for 5 minutes in an apparatus elevated to 120 cm from the floor, including two 35 cm length and 5cm wide open arms and two closed arms with same dimensions and surrounded with 15 cm high wall. The open field (OF) test was conducted in a 40 × 40 × 25 cm arena illuminated at 30 lux, where animals were allowed to freely explore for 10 minutes. The novelty suppressed feeding (NSF) test was performed after a 16-hour food deprivation period. Animals were placed in the open field arena, where a food pellet was positioned at the center. The latency to eat was recorded, with a maximum test duration of 10 minutes. The splash test (ST) was performed by spraying water onto the back of mice placed in a clean cage. The frequency of grooming behavior was recorded over a 5-minute period. The forced swim test (FST) was carried out for 6 minutes using a glass cylinder tank filled with water (26 ± 1 °C) in a manner that the mouse cannot reach the floor with its tail. Subsequently, the total immobility time during the last 5 minutes of the test was recorded. A 2-day recovery period between each test was respected. All videos were analyzed using the ANY-maze software (Stoelting Co.), automatically count on the software in a blinded setting.

### Metabolic tests

Body composition (fat mass and lean mass) was measured in conscious mice using a quantitative magnetic resonance system (EchoMRI, Bruker). Fasting blood glucose level, glucose tolerance in response to intraperitoneal (IP, ipGTT) glucose injection (2 g/kg), insulin sensitivity in response to IP insulin injection (ITT, 0.75 U/kg) were measured after 5-hour fasting from 9:00 am to 2:00 pm. Blood glucose level was measured from tail prick (Accu-check Performa glucometer; Roche) at 0 (fasting blood glucose), 15, 30, 45, 60, 90 and 120 minutes post-glucose (ipGTT) or -insulin (ITT) injection.

### Bulk RNA sequencing and analysis

RNAs were extracted as previously described (Martin et al., 2025) from hippocampi, prefrontal cortex or striatum of male C57BL/6J mice fed 25% fructose for 8 weeks using the Allprep DNA/RNA/ Protein Mini (Qiagen) according to manufacturer’s protocol. Bulk tissue RNA quality check and RNA-seq was performed by the University of Chicago Genomics Core. RNA concentration and purity were confirmed by Bioanalyzer 2100 (Agilent Technologies). All samples had a RIN score above 9.0. TruSeq stranded cDNA libraries were constructed using Oligo-dT mRNA method. RNA-seq was performed (approximately 50–60 million paired-end reads) using Illumina Nova-Seq 6000 at the Chicago Genomics Core facility.

Trimmed and paired-end reads only were aligned as previously described (Martin et al., 2025) using STAR two pass alignment indexed to the GRCm39/mmu 39 soft masked genome to identify splice or novel splice sites and re-indexed to quantify reads. Bioconductor packages in R Statistical Software (R version 4.4.1, R Core Team, 2024; https://www.bioconductor.org, v.3.19) were used for downstream read counts and differentially expressed genes (DEGs) analyses. Read counts were obtained with feature counts (Rsubread v2.16.1). Low-expressed genes were filtered (less than 10 reads in at least 3 samples). DESeq2 (v.1.44.0) was used for DEG analysis with GLM matrix weighted on experimental group.

#### Cell type-specific marker gene and pathway enrichment analyses

To identify which brain cell types were most transcriptionally affected by dietary fructose, we tested whether differentially expressed genes (DEGs) between fructose- and water-fed mice in the hippocampus, prefrontal cortex (PFC) or striatum were enriched for cell type-specific marker genes. Cell type marker gene sets were obtained from the BRETIGEA R package (markers_df_mouse_brain object) (McKenzie et al., 2018), which provides the top ∼1,000 marker genes for each of six major brain cell types — astrocytes, endothelial cells, microglia, neurons, oligodendrocytes, and oligodendrocyte precursor cells (OPCs; 430 markers) — derived from a meta-analysis of mouse brain cell type-specific gene expression data. All analyses were performed in Python (v3) using scipy.stats.fisher_exact and statsmodels.stats.multitest.multipletests. Pathway and process enrichment analysis of downregulated DEGs of the hippocampus was performed using Metascape (metascape.org) (Zhou et al., 2019). Default Metascape parameters were used unless otherwise specified (minimum count = 3, enrichment factor > 1.5, adjusted p < 0.01), and the top 10 enriched terms were visualized.

### Microglia isolation

Microglia isolation was performed as previously described (Herron et al., 2022). Briefly, female or male WT or GLUT5-KO mice (Coursan et al., 2025) fed standard chow or C57BL/6J mice fed fructose 25% or water were euthanized by intraperitoneal pentobarbital injection (Exagon®, 300 mg/kg), 30 min post-buprenorphine administration (0.1 mg/kg) before being transcardially perfused with ice-cold Hanks’ Balanced Salt Solution (HBSS, Gibco).

Following brain’s mechanical dissociation, single cell suspensions were prepared and centrifuged over a 37%/70% discontinuous Percoll gradient (GE Healthcare), mononuclear cells were isolated from the interphase. Isolated microglia were used for cell culture or RT-qPCR.

### Microglia cell culture

After isolation from the whole brain, microglia were filtered and plated at a density of 90,000–100,000 cells per well in 500 µL of A24943 SILAC Advanced DMEM/F-12 Flex culture medium (Fisher Scientifique, 15396278) containing 10% fetal bovine serum (Sigma, F9665) and 1% penicillin–streptomycin (Sigma, P4333-100mL) on poly-DL-ornithine-coated 48-well plates (Thermo Scientific Nunclon Delta Surface). The medium was supplemented with 0.7 mM L-arginine (Sigma, A5131-25G), 0.5 mM L-lysine (Sigma, L5626-100G), 2.5 mM D-glucose (Sigma, G8270), and 2.5 mM GlutaMAX supplement (Fisher, 11574466). To obtain M0 microglia (Butovsky et al., 2014), cytokines were added to the culture medium: 50 ng/mL TGF-β (Miltenyi Biotec, 130-095-066) and 10 ng/mL M-CSF (R&D Systems, 416-ML-010/CF). Microglia were treated with a physiological concentration of fructose (500 µM; Sigma, F0127). Cells were incubated for 5 days at 37°C in 5% CO₂. On day 5, the medium was replaced with cytokine-free medium for 24 hours. Cells were then treated with 100 ng/mL LPS (O127:B8, Sigma, L4516-1MG) for 3, 6, or 12 hours.

### Gene expression

Total RNAs from isolated microglia from hippocampi of control or fructose 25% fed C57BL/6J mice 3 hours post-LPS (250 µg/kg) treatment or saline; or primary cultured microglia were extracted using RNeasy Plus Mini Kit (Qiagen, 74106) according to the manufacturer’s protocol. RNA concentration and purity were determined using Nanodrop spectrophotometer (Nanodrop technologies). 10 µL of RNAs were reverse transcribed using Superscript IV VILO (Invitrogen, Life Technologies). cDNAs were amplified using commercially available FAM-labeled Taqman® probes (Applied Biosystems), [Glut5 (Mm00600311_m1), P2ry12 (Mm00446026_m1), Tgfbr2 (Mm03024091_m1), Tmem119 (Mm00525305_m1), Il-1b (Mm00434228_m1), Il-6 (Mm00446190_m1), Tnf-a (Mm00443258_m1), Tgf-β1 (Mm01178820_m1), B2m (Mm00437762_m1, used as reference gene)]. Real-time PCR reaction was performed in duplicate using a LightCycler® 480 instrument II (Roche). Data are normalized relative to B2m (2^(−ΔCt)).

### Immunohistochemistry

Iba1 and CD68 immunostaining were performed as previously described (Nakajima et al., 2026). Briefly, mice were euthanized by intraperitoneal pentobarbital injection (Exagon®, 300 mg/kg), 30 min post-buprenorphine administration (0.1 mg/kg) before being transcardially perfused with phosphate buffered saline (PBS) followed by 4 % paraformaldeide 24-48h after the last test performed. Dissected brains were post-fixed in 4 % paraformaldeide followed by 30 % sucrose, the brains were rapidly frozen in cold isopentane and stored at -80 °C until further processing. Coronal slices were obtained at 40 µm intervals with cryostat (Leica). After washing with PBS and blocking (1 h in 0.5 % Donkey serum, 0.1 % Triton, in PBS), slices were incubated with primary antibodies for anti-Iba1 (1:500, Rabbit, Fujifilm-WAKO 019-19741) and anti-CD68 (1:300, rat, Bio RAD MCA-1957) in the blocking solution overnight at 4 °C. Slices were washed with PBS and secondary antibodies were diluted in PBS (containing 5% donkey serum, 0.1% triton, 1:1000, anti-rabbit IgG Alexa 488, anti-rat IgG Alexa 594, Invitrogen) for 2 h at room temperature. Brain sections were mounted with vectashield which contains DAPI (Vector Labs) and imaged with Axio Imager A2 Zeiss fluorescent microscope (Carl Zeiss AG). Z-stack images were captured at 1 µm steps and stacked in ImageJ2/FIJI. First, the region of interest (ROI) was defined. Microglial cells within this ROI were then manually delineated to measure soma size. For CD68 quantification, a threshold was applied to quantify the area of CD68 signal into Iba1 signal.

### Statistical analysis

Statistical analysis was performed using GraphPad Prism 11 (GraphPad software). Data are expressed as mean ± SEM and individual values are plotted on graph. The number of animals is indicated by the individual data points on each bar. After normal Gaussian distribution was verified using the Kolmogorov-Smirnov test, appropriate parametric or non-parametric tests were chosen as mentioned in figure legends. All statistical tests used two-sided 0.05 significance threshold.

## RESULTS

### Dietary fructose-induced anxiety impacts microglia gene expression

Previous studies have reported that fructose consumption is associated with anxiety-like behavior and memory deficits in rodent models(Spagnuolo et al., 2020). To confirm these findings, adult male mice were fed with 25% fructose in their drinking water for 8 weeks (Figure 1A). As expected, fructose-fed mice exhibited increased body weight and fat mass and decreased lean mass (Supp. Figure 1B-D), along with increased fasting blood glucose level (Supp. Figure 1E), glucose intolerance (Supp. Figure 1F), and insulin resistance (Supp. Figure 1G). Behaviorally, male mice displayed anxiety-like behaviors in the EPM and the NFS tests (Figure 1C, D) as well as impaired self-care behaviors in the Splash test (Figure 1E) without any change in their locomotor activity (Figure 1B). In contrast, no depressive-like behavior was observed in the FST (Figure 1F). In addition, fructose-fed mice also showed memory deficits in the OLM test (Supp. Figure 1H) and the Morris water maze test without any change in the learning capacity (Supp. Figure 1I). Adult female mice exposed to a similar 8 weeks fructose diet (Supp. Figure 2A) showed modest changes in metabolic parameters, including increased body weight, fat mass, and fasting blood glucose level, but no significant impairments in glucose tolerance (Supp. Figure 2B-F). Interestingly, fructose-fed female mice did not exhibit anxiety or depressive-like behaviors in the EPM test, NSF test, or splash test (Supp. Figure 2H-J). In contrast, fructose-fed female mice displayed memory deficits in the OLM test (Supp. Figure 2K). Together, these results suggest that female mice are relatively protected from fructose-induced emotional disturbances but remain susceptible to memory impairments. Given the greater behavioral sensitivity observed in males, subsequent experiments were conducted primarily in male mice. Because anxiety can influence memory performance, we cannot conclude that fructose intake directly impairs memory. Therefore, we focused on elucidating the cellular mechanisms underlying fructose-induced emotional alterations.

**Figure 1:**
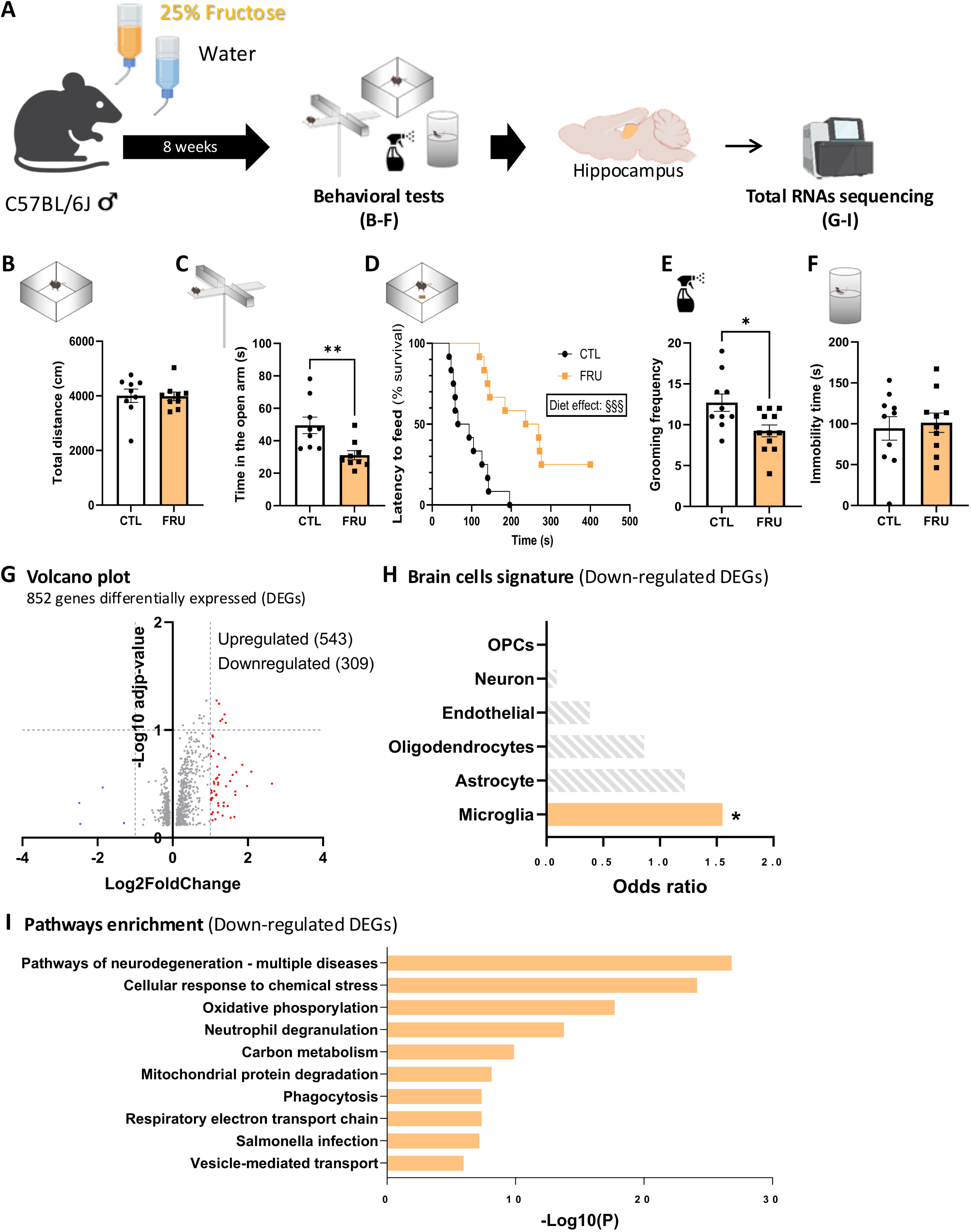
Fructose-induced emotional alterations is associated to transcriptional reprogramming in the hippocampus of adult male mice. (A) Experimental design: adult C57BL/6J male mice were exposed to 25% fructose in drinking water or control water (CTL) for 8 weeks before being subjected to a battery of emotional tests (B–F). Hippocampi were collected after behavioral assessments for bulk RNA sequencing (G–I). (B–F) Emotional behaviors. Total locomotion in the open field test (B), time spent in open arms of the elevated plus maze (C), latency to feed (% survival) in the novelty-suppressed feeding test (D), grooming frequency in the splash test (E), and immobility time in the forced swim test (F) in control (CTL) and fructose-fed (FRU) mice. (G–I) Hippocampal RNA-seq analysis. Volcano plot showing differentially expressed genes (DEGs; G), with the number of upregulated genes above 1 Log2FoldChange (Log2FC) in red and downregulated genes below -1 Log2FC indicated in blue. Cell type signature analysis obtained from the BRETIGEA R package (markers_df_mouse_brain object; McKenzie et al., 2018) of downregulated DEGs (H), highlighting enrichment for microglial gene sets. Enrichment analysis of the 10-most significant pathway of downregulated DEGs showing (I). Data are presented as mean ± SEM; *p < 0.05 *vs*. CTL (unpaired t-test).

**Figure 2:**
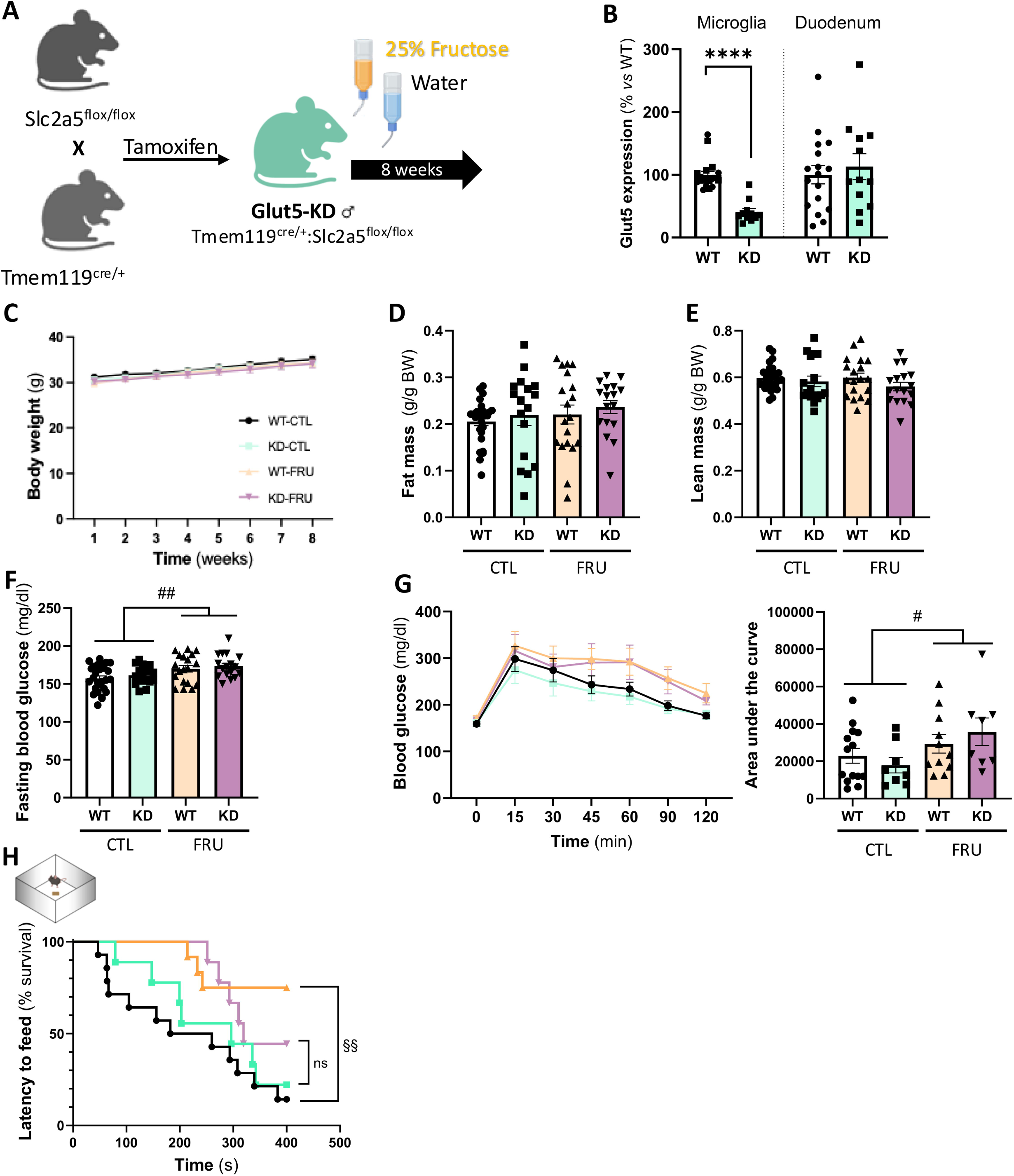
Microglial Glut5-knock down prevents fructose intake-induced anxiety but not metabolic disturbance. (A) Experimental design: Slc2a5flox/flox mice were crossed with Tmem119creERT2 mice to generate microglia-specific Glut5 knockdown (Glut5-KD; Tmem119cre/+:Slc2a5flox/flox and wild-type (WT) littermates (Slc2a5flox/flox cre negative). Adult male WT and Glut5-KD mice were treated with tamoxifen to induce Cre-mediated Glut5 deletion in microglia, then exposed to either control drinking water (CTL) or 25% fructose solution (FRU) for 8 weeks. (B) Validation of Glut5 knockdown. Glut5 expression in microglia and duodenum from WT and Glut5-KD mice. (C–G) Metabolic parameters. Body weight gain (C), fat mass (D), lean mass (E), 5-hour fasting blood glucose levels (F), and glucose tolerance during the intraperitoneal glucose tolerance test (ipGTT; G, left) with corresponding area under the curve (G, right) in WT and Glut5-KD mice under control (CTL) or FRU conditions. (H) Emotional behavior. Latency to feed (% survival) in the novelty-suppressed feeding test in WT and Glut5-KD mice under CTL or FRU conditions. Data are presented as mean ± SEM; p < 0.05, **** p < 0.0001 vs. WT; # p < 0.05, ## p < 0.01 vs. CTL (two-way ANOVA); $$ p < 0,01 vs. CTL (Logrank (Mantel-Cox) test).

To gain an unbiased and comprehensive understanding of the cellular and pathway-level impacts of dietary fructose, we performed bulk RNA sequencing on hippocampal, prefrontal cortex (PFC) and striatum tissues from fructose-fed male mice (Figure 1A). In the hippocampus, transcriptomic analysis identified 852 differentially expressed genes (DEGs) including 543 upregulated and 309 downregulated; of which 52 were upregulated with a Log2 fold change greater than 1, and 4 genes were downregulated with a Log2 fold change below -1 (Figure 1G). Interestingly, the hippocampus seemed particularly more sensitive than other brain areas since less genes are significantly differentially expressed in the prefrontal cortex (47 DEGs; Supp. Figure 3B) or the striatum (383 DEGs; Supp. Figure 3C). In the prefrontal cortex, only 10 genes were either upregulated with a Log2FC greater than 1, or downregulated with a Log2FC below -1 (Supp. Figure 3B) whereas no gene was upregulated with a Log2FC greater than 1, or downregulated with a Log2FC below -1 in the striatum (Supp. Figure 3C). Remarkably, Venn analysis illustrating the overlap of DEGs between hippocampus, PFC and striatum (Supp. Figure 3D), showed that fructose-induced DEGs were largely non-overlapping between hippocampus, PFC, and striatum, with most hippocampal DEGs being unique to this region, underscoring pronounced regional specificity of the transcriptional response. The stronger transcriptional response to fructose observed in the hippocampus is not associated with higher baseline expression of fructose transporters (Glut5, Glut8) or key fructose-metabolism enzymes (Supp. Figure 3E), suggesting that hippocampal vulnerability to fructose is not simply explained by increased expression of canonical fructose-metabolism genes.

**Figure 3:**
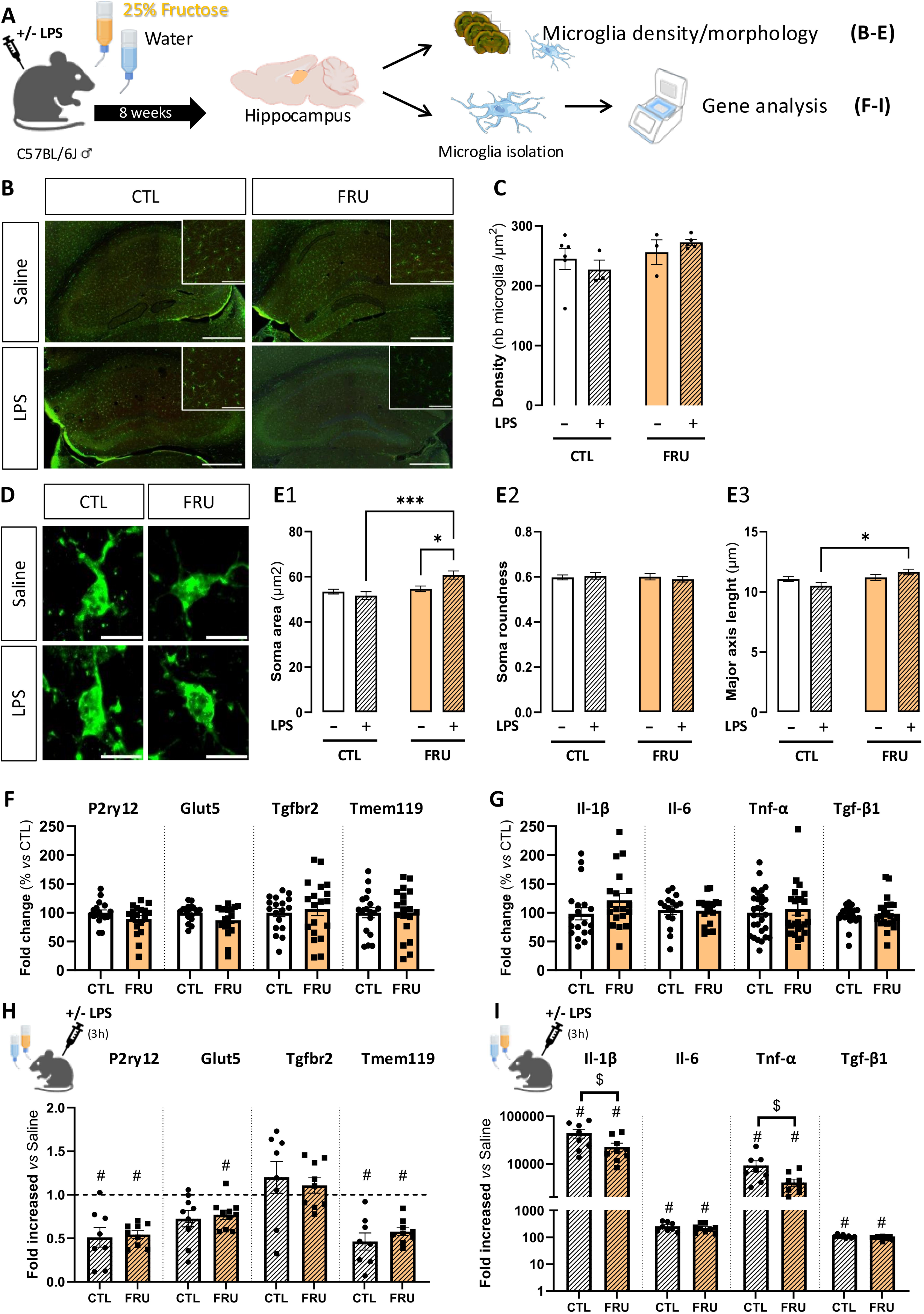
Fructose feeding alters microglia responsiveness. (A) Experimental design: adult. C57BL/6J male mice were exposed to 25% fructose in drinking water or control water (CTL) for 8 weeks before being treated with LPS (250 μg/kg). (B, C) Representative images (B, scale bar 500 µm; inset, scale bar 100 µm) and quantification (C) of microglia density in the hippocampus measured 3 hours post-LPS treatment. (D, E) Representative images (D, scale bar 10 µm) and quantification of microglia morphology measured 24 hours post-LPS treatment: soma area (E1), soma roundness (E2) and major axis length (E3). (F, G) Basal expression of the homeostatic genes P2ry12, Glut5, Tgfbr2 and Tmem119 (F) or cytokine genes Il1β, Il6, Tnfα and Tgfβ1 (G), expressed as fold change (%) relative to CTL. (H, I) Expression of the homeostatic genes P2ry12, Glut5, Tgfbr2 and Tmem119 (H) or cytokine genes Il1β, Il6, Tnfα and Tgfβ1 (I) measured 3 hours post-LPS treatment, expressed as fold increase vs. saline-treated mice under CTL or FRU conditions. Data are presented as mean ± SEM; * p < 0.05, *** p < 0.001 (two-way ANOVA followed by Tukey post-hoc test); # p < 0.05 vs. saline (unpaired t-test); $ p < 0.05 vs. CTL (two-way ANOVA followed by Tukey post-hoc test).

In the hippocampus, enrichment analysis using brain cell type marker genes obtained from the BRETIGEA R package (McKenzie et al., 2018) revealed that hippocampus downregulated DEGs were predominantly associated with a microglial signature, with no significant association from other brain cell types (Figure 1H), indicating that dietary fructose may primarily influence microglial functions. This observation is consistent with previous reports showing near-exclusive expression of Glut5 and a higher expression of Glut8 in microglia within the brain (Zhang et al., 2014). No significant cell-type association was detected for hippocampal upregulated DEGs (Supp. Figure 4A). Downregulated DEGs of the microglia signature are C1qa, C1qb, C1qc, Ctss, Hexb, Tyrobp. These genes of the C1q complement complex plus Hexb and Tyrobp genes are recognized homeostatic/surveillant microglia signature genes. Their reduction suggests that fructose feeding shifts hippocampal microglia away from this homeostatic state (i.e., altered microglial responsiveness/neuroimmune signaling), rather than broadly affecting neurons, glia, or vasculature. Moreover, a microglial signature was not observed among either upregulated or downregulated DEGs of the PFC or striatum (Supp. Figure 4B, C), confirming the particular vulnerability of the hippocampus microglia to fructose feeding. Finally, pathway enrichment analysis of downregulated hippocampal DEGs revealed significant suppression of pathways related to neurodegeneration, cellular responses to chemical stress, oxidative phosphorylation and mitochondrial protein degradation, as well as carbon metabolism and phagocytosis (Figure 1I). These changes suggest that chronic fructose intake impairs microglial metabolic and bioenergetic functions while weakening stress-response and clearance pathways in the hippocampus.

**Figure 4:**
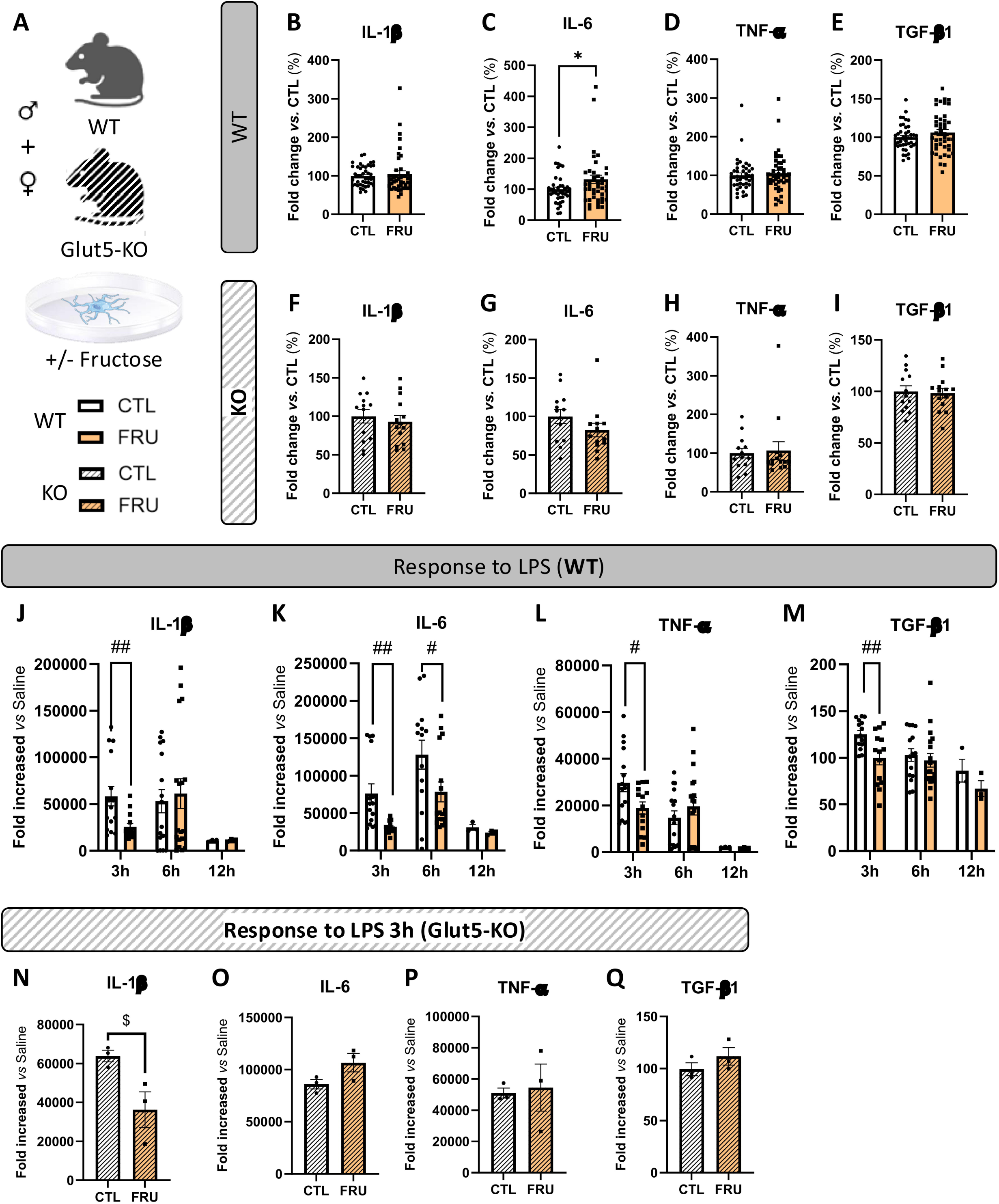
Fructose delays microglia response to LPS in a Glut5-dependent manner. **(A)** Experimental design: primary microglia were isolated from adult wild-type (WT) or whole-body Glut5 knockout (Glut5-KO) male and female mice and cultured with (FRU) or without (CTL) fructose (500 µM) for 5 days before being treated with LPS (100 ng/mL). (B–I) Basal cytokine expression in microglia from WT (B–E) or Glut5-KO (F–I): IL-1β (B, F), IL-6 (C, G), TNF-α (D, H), and TGF-β1 (E, I), expressed as fold change (%) relative to CTL. (J–M) Time course of cytokine induction in microglia cultured from WT mice following LPS stimulation: IL-1β (J), IL-6 (K), TNF-α (L), and TGF-β1 (M), represented as fold increase *vs.* saline-treated cells at 3, 6, and 12 h in the presence (FRU) or absence (CTL) of fructose. (N–Q) Cytokine induction at 3 h after LPS treatment in microglia cultured from Glut5-KO mice: IL-1β (N), IL-6 (O), TNF-α (P), and TGF-β1 (Q), expressed as fold increase *vs.* saline-treated cells under CTL or FRU conditions. Data are presented as mean ± SEM; *p < 0.05 *vs.* CTL (unpaired t-test); # p < 0.05, ## p < 0.01 *vs.* CTL (two-way ANOVA repeated measures followed by Bonferoni post-hoc test); $ p < 0.05 *vs.* CTL (two-way ANOVA followed by Tukey post-hoc test).

### Fructose-feeding induced anxiety is dependent on fructose uptake by microglia

Since RNA-sequencing analyses of the hippocampus revealed that fructose feeding impacts cell signature representative of microglia and that the main fructose transporter Glut5 is almost exclusively expressed in this cell type, we knocked down Glut5 selectively in microglia (Glut5-KD mice). Therefore, we created the Glut5flox/flox mice that we crossed with Tmem119-creERT2 mice (Figure 2A). After tamoxifen treatment, Glut5 expression was significantly decreased by 59.3 ± 5.3 % selectively in microglia and not in the proximal part of the gut (Figure 2B) were its expression is the most abundant (Douard and Ferraris, 2013). Glut5-KD mice were fed 25% fructose in their drinking water for 8 weeks as shown above. Fructose feeding did not alter total body weight, fat mass or lean mass both in WT and Glut5-KD mice (Figure 2C-E). In contrast, as expected, fructose feeding increased fasting blood glucose level and induced glucose intolerance in both WT and Glut5-KD mice (Figure 2F, G). Surprisingly, fructose feeding failed to decrease the time spent in the open arm of the EPM or the frequency of grooming in the Splash test in WT or Glut5-KD mice (Supp. Figure 5B, C). Nevertheless, in the novelty-suppressed feeding test, fructose feeding increased the latency to feed in WT mice but not in Glut5-KD mice (Figure 2H), suggesting that the knockdown of Glut5 selectively in microglia prevented fructose-induced anxiety.

### Fructose feeding impairs microglia morphology and responsiveness

Transcriptomic analyses suggested that fructose feeding affects pathways related to microglial function and neuroimmune signaling. Therefore, we assessed microglial density, morphology, and gene expression in the hippocampus of mice exposed to 25% fructose in drinking water. These parameters were examined under basal conditions and following lipopolysaccharide (LPS) challenge to evaluate microglial responsiveness and cytokine expression (Figure 3A). Under basal conditions, fructose feeding did not alter microglial density (Figure 3B, C), morphology (Figure 3D, E), or the expression of microglial homeostatic markers and cytokines, except for a modest increase in Il6 expression (Figure 3F, G). In contrast, LPS challenge revealed altered microglial responsiveness in fructose-fed mice. Although LPS did not modify microglial density (Figure 3B, C), microglia from fructose-fed mice displayed a larger soma area (Figure 3D, 3E1) and increased major-axis length compared with LPS-treated control mice (Figure 3D, 3E3), without changes in soma roundness (Figure 3D, 3E2). LPS reduced the expression of the homeostatic markers P2ry12, Glut5, Tgfbr2, and Tmem119 in both groups (Figure 3H). However, microglia isolated from fructose-fed mice showed a blunted cytokine response to LPS, with Il-1β and Tnf-α expression reduced by approximately 50% compared with microglia from LPS-treated control mice (Figure 3I; Il-1β: 44083 ± 9236 vs. 22894 ± 4499; Tnf-α: 9306 ± 2312 vs. 4054 ± 736, respectively), whereas Il-6 and Tgf-β1 induction was not significantly affected (Figure 3I). Wang et al. recently described that perinatal fructose exposure impairs microglia phagocytosis (Wang et al., 2025). We therefore assessed CD68 immunoreactivity within Iba1-positive cells as an index of microglial lysosomal/phagolysosomal activity. Under basal conditions, CD68 immunoreactivity was reduced in fructose-fed animals (Supp. Figure 6B, C), in line with our RNAseq pathway analysis indicating altered phagocytosis-related signalling (Figure 1I). Following LPS challenge, CD68 immunoreactivity was not significantly altered in control animals, whereas it was increased in fructose-fed animals (Supp. Figure 6B, C). These data suggest that fructose feeding impairs basal microglial phagolysosomal content without compromising the phagolysosomal response to an inflammatory challenge. Together, these findings indicate that fructose feeding does not induce overt basal microglial alterations apart from a reduction in phagolysosomal activity, but modifies microglial morphological and transcriptional responsiveness to inflammatory challenge.

### Fructose directly impairs microglia responsiveness *in vitro*

To assess whether fructose directly modulates cytokine production in vitro, primary adult microglia isolated from wild-type (WT) and Glut5 knockout (Glut5-KO) mice were exposed to 500 µM fructose and subsequently challenged with LPS (Figure 4A). Under basal conditions, fructose exposure did not alter the expression of homeostatic genes P2ry12, Tgfbr2 in either WT or Glut5-KO microglia (Supp. Figure 7A-F). Fructose induced a slight increase in IL-6 expression in WT microglia, but not in Glut5-KO cells (Figure 4C, G). In contrast, expression levels of other pro-inflammatory cytokines (IL-1β, Tnf-α) as well as the anti-inflammatory cytokine Tgf-β1 remained unchanged in WT or Glut5-KO microglia (Figure 4B-I). In microglia isolated from WT mice, LPS increased the expression IL-1β, IL-6, Tnf-α and Tgf-β1 at 3, 6 or 12h post-treatment (Figure 4J-M). Notably, fructose exposure decreased expression levels of IL-6, IL-1β, Tnf-α, and Tgf-β1 at 3 hours post-LPS stimulation, but this effect was absent at 6 and 12 hours, except for sustained Il-6 suppression at 6 hours (Figure 4J-M). In Glut5-KO microglia, cytokine expression 3 hours post-LPS was unaffected, except for a change in IL-1β (Figure 4N-Q), indicating that fructose’s modulatory effect is largely GLUT5-dependent. Collectively, these results indicate that fructose directly act on microglia through Glut5 to impair their responsiveness to inflammatory challenge.

## DISCUSSION

Several animal studies have suggested that excessive fructose intake induce brain health dysfunction, including anxiety-like behaviors, depressive-like symptoms, and cognitive deficits(Spagnuolo et al., 2020). The aim of the present study was to elucidate the cellular and molecular mechanisms underlying these neurobehavioral effects. Our findings identify microglia as a key mediator. An unbiased transcriptomic approach (bulk RNA-seq) revealed that fructose feeding predominantly alters hippocampal genes enriched for a microglial signature. Selective knockdown of the fructose transporter Glut5 in microglia prevented the development of fructose-induced anxiety-like behavior. Finally, our data indicate that fructose impairs microglial responsiveness to promote anxiety.

### Fructose feeding impairs brain health

In our study, 25% fructose in the drinking water for 8 weeks induced anxietylike, but not depressivelike, behaviors in adult male mice, whereas adult females were largely spared these emotional alterations. These findings are in line with some, but not all, rodent studies investigating the behavioral consequences of high fructose intake. Discrepancies across the literature likely reflect important differences in experimental design, including sex, species or strain (mouse *vs.* rat), fructose concentration, duration and mode of administration (drinking water *vs.* chow), and the developmental period during which the diet is applied, all of which can strongly influence the behavioral phenotypes (Spagnuolo et al., 2020; De Souza et al., 2021). In our conditions, adult female mice remained largely protected from fructoseinduced anxietylike behavior, despite some metabolic alterations. One may hypothesize that extending the duration of fructose exposure would eventually lead to the emergence of anxietylike behaviors in females. This view is supported by studies in female mice fed obesogenic high-fat diets, where emotional phenotypes emerge only after prolonged exposures: for example, anxiodepressive–like behaviors appear after 11–24 weeks of high-fat feeding in females, whereas shorter protocols can be behaviorally neutral or even anxiolytic (Krishna et al., 2016; Décarie-Spain et al., 2021; Huang et al., 2024). These studies support the notion that in females, a longer duration of fructose or obesogenic feeding may be required to unmask anxietylike behaviors.

Consistent with several data from the literature(Franco-Pérez, 2024), we show that 25% fructose feeding induces memory deficits in both adult male and female mice. Nevertheless, heightened anxiety is known to impair hippocampaldependent memory performance in rodents (Moreira et al., 2016), and pharmacological attenuation of stress hormones or stress-related signaling can normalize memory performance in stressed animals (Béracochéa et al., 2011). Thus, the memory deficits observed after fructose feeding in male cannot be unequivocally attributed to a direct effect on cognitive processes and may instead reflect, at least in part, secondary consequences of anxiety. In female mice, by contrast, the memory effect of fructose may more likely reflect a direct effect on cognition, since none of the behavioral tests performed detected increased anxiety.

### Fructose-induced anxiety in microglia-dependent

Several *in vivo* and *in vitro* studies have reported that fructose affects microglia (Ting, 2024b). However, these studies generally tested predefined hypotheses and targeted a limited set of markers or pathways. Here, using an unbiased RNA-seq approach, we show that fructose feeding alters gene expression patterns reminiscent of a microglial signature. These data support the idea that microglia are a direct target of dietary fructose. This is consistent with the fact that, among brain cells, Glut5 is expressed predominantly in microglia (Zhang et al., 2014). Importantly, and in line with a recent study in a model of perinatal fructose exposure (Wang et al., 2025), we show that the selective knockdown of Glut5 in microglia prevents the development of fructose-induced anxiety. Although this strategy does not generate a complete knockout, microglial Glut5 expression was reduced by approximately 60%, which is consistent with the partial recombination efficiencies typically obtained with Tmem119-CreER-based models after tamoxifen induction. Notably, this level of knockdown was sufficient to prevent the emergence of anxiety-like behavior in the NSF test, indicating that partial reduction of microglial Glut5 is enough to block this phenotype. Thus, fructose failed to induce anxiety-like behavior in Glut5-KD mice in this assay. This finding indicates that microglial fructose uptake via Glut5 is a necessary step in the development of anxiety. By contrast, this protective effect of Glut5 knockdown could not be evaluated in the EPM or splash tests, because fructose did not induce an anxiety-like phenotype in WT animals in these assays. Notably, Glut5-KD and WT littermates were generated on a C57BL/6N background, whereas the naïve mice used in the rest of the study were on a C57BL/6J background. C57BL/6J and C57BL/6N mice differ behaviorally, underscoring the influence of genetic background on emotional phenotypes (Bryant et al., 2008). These substrain-specific differences also extend to responses to metabolic challenges, as C57BL/6J and C57BL/6N mice exhibit divergent susceptibility to diet-induced metabolic and anxiety-related phenotypes (Maroni et al., 2020). This variability should therefore be taken into account when interpreting fructose-induced behavioral outcomes and underscores the need to use multiple behavioral assays when assessing emotional phenotypes.

### Fructose alters microglia responsiveness

Our study shows that microglia are a key mediator of fructose-induced anxiety. An important question is how fructose alters microglial function to promote anxiety. Several studies have reported that fructose feeding increases pro-inflammatory cytokine expression in distinct brain regions, including the hippocampus (Cigliano et al., 2018; Li et al., 2019; Fierros-Campuzano et al., 2022). However, these studies did not demonstrate that these changes specifically originated from microglia. In addition, in vitro studies have suggested that fructose can directly increase cytokine expression in microglia, but most of these experiments used supraphysiological fructose concentrations (> 5 mM) (Xu et al., 2016; Cigliano et al., 2018; Mizuno et al., 2021; Ting, 2024b), far above estimated brain fructose concentrations, which are approximately 0.3 mM (Hwang et al., 2015, 2017). In our study, physiological fructose exposure did not significantly increase cytokine gene expression, although IL-6 showed a modest increase. This discrepancy could be due to the level of fructose used. Supraphysiologic concentration of fructose could stress microglia which may produce cytokines in response. By contrast, in our study, we found that fructose blunted the microglial cytokine response to LPS *in vitro* in microglia culture as well as *in vivo* in fructose fed animals treated with LPS. These data suggest that rather than promoting neuroinflammation, fructose impairs microglia reactivity. This reduced responsiveness, is consistent with a recent study showing that perinatal fructose exposure impairs microglial phagocytic function (Wang et al., 2025), although in a developmental rather than adult paradigm. In line with this, we found that CD68 immunoreactivity within Iba1-positive cells was significantly decreased by fructose feeding under basal conditions. Following LPS challenge, CD68 immunoreactivity was unchanged in control animals but increased in fructose-fed animals, indicating that the microglial response to LPS is impaired at the level of CD68 as well as cytokine expression. Taken together, these data suggest that fructose does not primarily drive microglia toward an activated inflammatory state, but instead alters their functional responsiveness.

Importantly, our RNA-seq data also point to alterations in intracellular metabolic pathways in the hippocampus of fructose-fed mice. This observation is consistent with recent data showing that fructose can reprogram microglial metabolism and alter their activation state (Ting, 2024b), and raises the possibility that fructose reprograms microglial metabolism, thereby limiting the energetic flexibility required for appropriate activation and plasticity. Such metabolic impairment could contribute to defective microglial surveillance and reduced capacity to adapt to environmental or inflammatory challenges. Altogether, these findings support the view that chronic fructose exposure promotes a maladaptive microglial state that may contribute to anxiety-related circuit dysfunction. One question that remains is how impaired microglial responsiveness may contribute to anxiety. A growing body of evidence indicates that microglial dysfunction can alter anxiety-related behavior, and both loss of specific microglial populations and pharmacological or genetic disruption of microglial function have been associated with anxiety-like phenotypes (Lehmann et al., 2019; Cordeiro et al., 2024). Because microglia regulate synaptic pruning, trophic support, and inflammatory signaling, a hypo-reactive microglial state may impair circuit remodeling and weaken the homeostatic control of networks implicated in anxiety including the hippocampus (Wang et al., 2022).

### Conclusion

Our results support a direct contribution of microglial dysfunction to fructose-induced anxiety. However, the gut–brain axis is increasingly recognized as a key regulator of emotional behavior (Cryan et al., 2020). Fructose feeding also alters the gut microbiota composition and metabolism (Guney et al., 2023), and our recent work showed that fructose malabsorption–induced dysbiosis is associated with anxiety in humans and animal models (Coursan et al., 2025). Therefore, fructose-induced anxiety likely reflects the combined action of gut-derived and microglia-dependent pathways. In this context, microglia emerge as a central, but not exclusive, mediator of the behavioral effects of fructose. Overall, our study identifies microglia as a critical mediator of fructose-induced anxiety, while leaving open the possibility that gut dysbiosis acts in parallel to shape the full behavioral phenotype. From a nutritional perspective, our study further supports the idea that excessive fructose intake can directly affect the brain and contribute to mental health disorders. Although the fructose concentration used in our in vivo experiments was relatively high, it remains within the range of intake reported in some human populations. Our findings therefore reinforce current public health recommendations, including the WHO guidance to reduce free sugar intake to below 5% of total energy intake.

## DECLARATIONS

### Ethics approval and consent to participate

All rodent experimental procedures were conducted in accordance with the European directive 2010/63/UE and approved by the French Ministry of Research and local ethics committees (APAFIS# 19870 and APAFIS# 13272).

### Consent for publication

Not applicable.

## Funding

This work was supported by grants from the Société Française de Nutrition (SFN, XF), the Fondation pour la Recherche Médicale (FRM, XF, VD), the Fondation pour la Recherche sur le Cerveau (FRC, XF, VD), the département AlimH INRAE (XF, AC), Bordeaux INP (AC), the ANR (Micro2Sweet, XF), the Bordeaux Neurocampus (XF) and the Région Nouvelle Aquitaine (CHESS Exomarquage, SL).

### Availability of data and materials

All data are included in this article and its supplementary information files.

### Authors’ contributions

XF designed and supervised the project and wrote the manuscript. AC, TD, MPDL performed cell and rodent behavior experiments. MH, SES performed RNAseq analyses. CM, AC performed microglia analysis. All authors edited/reviewed the manuscript.

## Acknowledgments

XF and AC are grateful to Bordeaux INP and INRAE for their support. Authors are thankful to the following: (i) the staff of the CIRCE (Behavioral Engineering Centre) facility of Bordeaux Neurocampus and the staff of the Service Commun des Animaleries (SCA) of the Université de Bordeaux for animal care, housing and breeding and technical support throughout this study, (ii) the CRYME core facility of Bordeaux Neurocampus for their support in the generation and breeding of the Glut5-KD mouse line, (iii) the RRI Food4BrainHealth for the inspiring discussions of the data.

## Declaration of Generative AI and AI-assisted Technologies in the Writing Process

During the preparation of this manuscript, the authors used Claude AI to assist with English-language editing and improvement of readability. No scientific interpretation, data analysis was delegated to the tool. The authors reviewed and edited all suggestions and take full responsibility for the content of the manuscript.

## REFERENCES

Barma, M. D., B. M. Purohit, H. Priya, S. Malhotra, U. S. Bhadauria, and R. Duggal, 2026, Sweet Misery: Association of Sugar Consumption With Anxiety and Depression—A Systematic Review: Obesity Reviews, v. 27, no. 1, p. e70003, doi:10.1111/obr.70003.

Béracochéa, D., C. Tronche, M. Coutan, R. Dorey, F. Chauveau, and C. Piérard, 2011, Interaction between Diazepam and Hippocampal Corticosterone after Acute Stress: Impact on Memory in Middle-Aged Mice: Frontiers in Behavioral Neuroscience, v. 5, p. 14, doi:10.3389/fnbeh.2011.00014.

Bryant, C. D., N. N. Zhang, G. Sokoloff, M. S. Fanselow, H. S. Ennes, A. A. Palmer, and J. A. McRoberts, 2008, Behavioral Differences among C57BL/6 Substrains: Implications for Transgenic and Knockout Studies: Journal of neurogenetics, v. 22, no. 4, p. 315–331, doi:10.1080/01677060802357388.

Butovsky, O. et al., 2014, Identification of a unique TGF-β-dependent molecular and functional signature in microglia: Nature Neuroscience, v. 17, no. 1, p. 131–143, doi:10.1038/nn.3599.

Cigliano, L., M. S. Spagnuolo, R. Crescenzo, R. Cancelliere, L. Iannotta, A. Mazzoli, G. Liverini, and S. Iossa, 2018, Short-Term Fructose Feeding Induces Inflammation and Oxidative Stress in the Hippocampus of Young and Adult Rats: Molecular Neurobiology, v. 55, no. 4, p. 2869– 2883, doi:10.1007/s12035-017-0518-2.

Colonna, M., and O. Butovsky, 2017, Microglia Function in the Central Nervous System During Health and Neurodegeneration: Annual Review of Immunology, v. 35, p. 441–468, doi:10.1146/annurev-immunol-051116-052358.

Cordeiro, R. C., G. Scaini, and J. Quevedo, 2024, Are microglia in charge of controlling stress-response behavior? Molecular Psychiatry, v. 29, no. 8, p. 2599–2600, doi:10.1038/s41380-024-02511-x.

Coursan, A. et al., 2025, Fructose malabsorption induces dysbiosis and increases anxiety in male human and animal models: Brain, Behavior, and Immunity, p. 106221, doi:10.1016/j.bbi.2025.106221.

Cryan, J. F., K. J. O’Riordan, K. Sandhu, V. Peterson, and T. G. Dinan, 2020, The gut microbiome in neurological disorders: The Lancet. Neurology, v. 19, no. 2, p. 179–194, doi:10.1016/S1474-4422(19)30356-4.

De Souza, L. et al., 2021, Impact of different fructose concentrations on metabolic and behavioral parameters of male and female mice: Physiology & Behavior, v. 228, p. 113187, doi:10.1016/j.physbeh.2020.113187.

Décarie-Spain, L., C. Hryhorczuk, D. Lau, É. Jacob-Brassard, A. Fisette, and S. Fulton, 2021, Prolonged saturated, but not monounsaturated, high-fat feeding provokes anxiodepressive-like behaviors in female mice despite similar metabolic consequences: Brain, Behavior, & Immunity -Health, v. 16, p. 100324, doi:10.1016/j.bbih.2021.100324.

Douard, V., and R. P. Ferraris, 2013, The role of fructose transporters in diseases linked to excessive fructose intake: The Journal of Physiology, v. 591, no. 2, p. 401–414, doi:10.1113/jphysiol.2011.215731.

Fierros-Campuzano, J., P. Ballesteros-Zebadúa, J. Manjarrez-Marmolejo, P. Aguilera, M. Méndez-Diaz, O. Prospero-García, and J. Franco-Pérez, 2022, Irreversible hippocampal changes induced by high fructose diet in rats: Nutritional Neuroscience, v. 25, no. 6, p. 1325– 1337, doi:10.1080/1028415X.2020.1853418.

Franco-Pérez, J., 2024, Mechanisms Underlying Memory Impairment Induced by Fructose: Neuroscience, v. 548, p. 27–38, doi:10.1016/j.neuroscience.2024.04.001.

Fulton, S., L. Décarie-Spain, X. Fioramonti, B. Guiard, and S. Nakajima, 2022, The menace of obesity to depression and anxiety prevalence: Trends in endocrinology and metabolism: TEM, v. 33, no. 1, p. 18–35, doi:10.1016/j.tem.2021.10.005.

Gillespie, K. M., M. J. White, E. Kemps, H. Moore, A. Dymond, and S. E. Bartlett, 2023, The Impact of Free and Added Sugars on Cognitive Function: A Systematic Review and Meta-Analysis: Nutrients, v. 16, no. 1, p. 75, doi:10.3390/nu16010075.

Guney, C., N. B. Bal, and F. Akar, 2023, The impact of dietary fructose on gut permeability, microbiota, abdominal adiposity, insulin signaling and reproductive function: Heliyon, v. 9, no. 8, p. e18896, doi:10.1016/j.heliyon.2023.e18896.

Herron, S., J.-C. Delpech, C. Madore, and T. Ikezu, 2022, Using mechanical homogenization to isolate microglia from mouse brain tissue to preserve transcriptomic integrity: STAR protocols, v. 3, no. 4, p. 101670, doi:10.1016/j.xpro.2022.101670.

Huang, H., J. Huang, W. Lu, Y. Huang, R. Luo, L. Bathalian, M. Chen, and X. Wang, 2024, A Four-Week High-Fat Diet Induces Anxiolytic-like Behaviors through Mature BDNF in the mPFC of Mice: Brain Sciences, v. 14, no. 4, p. 389, doi:10.3390/brainsci14040389.

Hwang, J. J., L. Jiang, M. Hamza, F. Dai, R. Belfort-DeAguiar, G. Cline, D. L. Rothman, G. Mason, and R. S. Sherwin, 2017, The human brain produces fructose from glucose: JCI insight, v. 2, no. 4, p. e90508, doi:10.1172/jci.insight.90508.

Hwang, J. J., A. Johnson, G. Cline, R. Belfort-DeAguiar, D. Snegovskikh, B. Khokhar, C. S. Han, and R. S. Sherwin, 2015, Fructose levels are markedly elevated in cerebrospinal fluid compared to plasma in pregnant women: PloS One, v. 10, no. 6, p. e0128582, doi:10.1371/journal.pone.0128582.

Krishna, S., Z. Lin, C. B. de La Serre, J. J. Wagner, D. H. Harn, L. M. Pepples, D. M. Djani, M. T. Weber, L. Srivastava, and N. M. Filipov, 2016, Time-dependent behavioral, neurochemical, and metabolic dysregulation in female C57BL/6 mice caused by chronic high-fat diet intake: Physiology & behavior, v. 157, p. 196–208, doi:10.1016/j.physbeh.2016.02.007.

Lehmann, M. L., T. K. Weigel, C. N. Poffenberger, and M. Herkenham, 2019, The Behavioral Sequelae of Social Defeat Require Microglia and Are Driven by Oxidative Stress in Mice: Journal of Neuroscience, v. 39, no. 28, p. 5594–5605, doi:10.1523/JNEUROSCI.0184-19.2019.

Li, J.-M., R. Yu, L.-P. Zhang, S.-Y. Wen, S.-J. Wang, X.-Y. Zhang, Q. Xu, and L.-D. Kong, 2019, Dietary fructose-induced gut dysbiosis promotes mouse hippocampal neuroinflammation: a benefit of short-chain fatty acids: Microbiome, v. 7, no. 1, p. 98, doi:10.1186/s40168-019-0713-7.

Maroni, M. J. et al., 2020, Substrain specific behavioral responses in male C57BL/6N and C57BL/6J mice to a shortened 21-hour day and high-fat diet: Chronobiology International, v. 37, no. 6, p. 809–823, doi:10.1080/07420528.2020.1756840.

Martin, H. et al., 2024, Insulin modulates emotional behavior through a serotonin-dependent mechanism: Molecular Psychiatry, v. 29, no. 6, p. 1610–1619, doi:10.1038/s41380-022-01812-3.

Martin, M. et al., 2025, Preventing effect of plant extracts and omega-3 on age-related cognitive decline in male mice: Brain, Behavior, & Immunity - Health, v. 49, p. 101127, doi:10.1016/j.bbih.2025.101127.

Martin, H., S. Bullich, B. P. Guiard, and X. Fioramonti, 2021, The impact of insulin on the serotonergic system and consequences on diabetes-associated mood disorders: Journal of Neuroendocrinology, v. 33, no. 4, p. e12928, doi:10.1111/jne.12928.

McKenzie, A. T. et al., 2018, Brain Cell Type Specific Gene Expression and Co-expression Network Architectures: Scientific Reports, v. 8, no. 1, p. 8868, doi:10.1038/s41598-018-27293-5.

Mizuno, T. M., P. S. Lew, and G. Jhanji, 2021, Regulation of the Fructose Transporter Gene Slc2a5 Expression by Glucose in Cultured Microglial Cells: International Journal of Molecular Sciences, v. 22, no. 23, p. 12668, doi:10.3390/ijms222312668.

Moreira, P. S., P. R. Almeida, H. Leite-Almeida, N. Sousa, and P. Costa, 2016, Impact of Chronic Stress Protocols in Learning and Memory in Rodents: Systematic Review and Meta-Analysis: PLoS ONE, v. 11, no. 9, p. e0163245, doi:10.1371/journal.pone.0163245.

Nakajima, S., A. Kabahizi, S. Aubailly, S. Naili-Douaouda, A. Tomita, A. Bosson, T. Alquier, C. Murphy-Royal, and S. Fulton, 2026, Anxiety induced by systemic inflammation involves microglial engulfing of nucleus accumbens inputs and diminished excitability of dopamine D1 receptor neurons: Brain, Behavior, and Immunity, v. 131, p. 106154, doi:10.1016/j.bbi.2025.106154.

Spagnuolo, M. S., S. Iossa, and L. Cigliano, 2020, Sweet but Bitter: Focus on Fructose Impact on Brain Function in Rodent Models: Nutrients, v. 13, no. 1, p. E1, doi:10.3390/nu13010001.

Tappy, L. et al., 2018, French Recommendations for Sugar Intake in Adults: A Novel Approach Chosen by ANSES: Nutrients, v. 10, no. 8, p. 989, doi:10.3390/nu10080989.

Tappy, L., and K.-A. Lê, 2010, Metabolic effects of fructose and the worldwide increase in obesity: Physiological Reviews, v. 90, no. 1, p. 23–46, doi:10.1152/physrev.00019.2009.

Ting, K. K. Y., 2024a, Fructose overconsumption-induced reprogramming of microglia metabolism and function: Frontiers in Immunology, v. 15, p. 1375453, doi:10.3389/fimmu.2024.1375453.

Ting, K. K. Y., 2024b, Fructose overconsumption-induced reprogramming of microglia metabolism and function: Frontiers in Immunology, v. 15, p. 1375453, doi:10.3389/fimmu.2024.1375453.

Wang, Z. et al., 2025, Early life high fructose impairs microglial phagocytosis and neurodevelopment: Nature, v. 644, no. 8077, p. 759–768, doi:10.1038/s41586-025-09098-5.

Wang, H., Y. He, Z. Sun, S. Ren, M. Liu, G. Wang, and J. Yang, 2022, Microglia in depression: an overview of microglia in the pathogenesis and treatment of depression: Journal of Neuroinflammation, v. 19, no. 1, p. 132, doi:10.1186/s12974-022-02492-0.

Xu, M.-X., R. Yu, L.-F. Shao, Y.-X. Zhang, C.-X. Ge, X.-M. Liu, W.-Y. Wu, J.-M. Li, and L.-D. Kong, 2016, Up-regulated fractalkine (FKN) and its receptor CX3CR1 are involved in fructose-induced neuroinflammation: Suppression by curcumin: Brain, Behavior, and Immunity, v. 58, p. 69–81, doi:10.1016/j.bbi.2016.01.001.

Zemdegs, J. et al., 2019, Metformin Promotes Anxiolytic and Antidepressant-Like Responses in Insulin-Resistant Mice by Decreasing Circulating Branched-Chain Amino Acids: The Journal of Neuroscience: The Official Journal of the Society for Neuroscience, v. 39, no. 30, p. 5935– 5948, doi:10.1523/JNEUROSCI.2904-18.2019.

Zemdegs, J., G. Quesseveur, D. Jarriault, L. Pénicaud, X. Fioramonti, and B. P. Guiard, 2016, High-fat diet-induced metabolic disorders impairs 5-HT function and anxiety-like behavior in mice: British Journal of Pharmacology, v. 173, no. 13, p. 2095–2110, doi:10.1111/bph.13343.

Zhang, Y. et al., 2014, An RNA-sequencing transcriptome and splicing database of glia, neurons, and vascular cells of the cerebral cortex: The Journal of Neuroscience: The Official Journal of the Society for Neuroscience, v. 34, no. 36, p. 11929–11947, doi:10.1523/JNEUROSCI.1860-14.2014.

Zhou, Y., B. Zhou, L. Pache, M. Chang, A. H. Khodabakhshi, O. Tanaseichuk, C. Benner, and S. K. Chanda, 2019, Metascape provides a biologist-oriented resource for the analysis of systems-level datasets: Nature Communications, v. 10, no. 1, p. 1523, doi:10.1038/s41467-019-09234-6.

