## Supplemental Figures for "Dietary fructose promotes anxiety through impairment of microglia physiology"

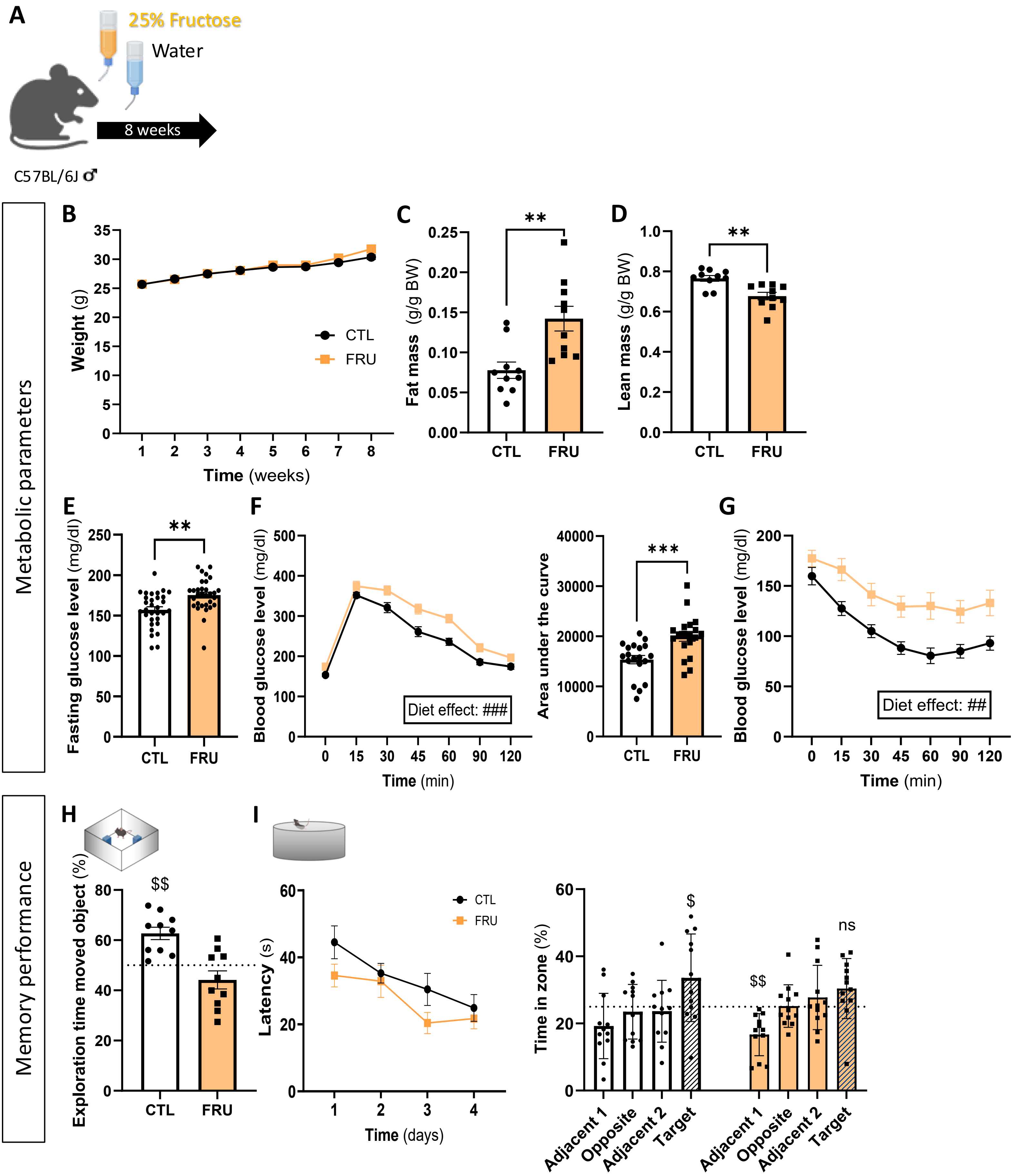
A
25% Fructose
Water
8 weeks
C57BL/6J ♂
Metabolic parameters
B
Weight (g)
Time (weeks)
CTL
FRU
C
Fat mass (g/g BW)
**
D
Lean mass (g/g BW)
**
E
Fasting glucose level (mg/dl)
**
F
Blood glucose level (mg/dl)
Diet effect: ###
Time (min)
Area under the curve
***
G
Blood glucose level (mg/dl)
Diet effect: ##
Time (min)
Memory performance
H
Exploration time moved object (%)
$$
I
Latency (s)
Time (days)
Time in zone (%)
$
$$
ns
Adjacent 1
Opposite
Adjacent 2
Target

**<u>Supp. Figure 1:</u> Metabolic parameters and memory performance of chronic fructose intake in adult male mice.** (A) Experimental design: adult C57BL/6J male mice were exposed to 25% fructose in drinking water or control water for 8 weeks. (B–G) Metabolic parameters. Body weight gain (B), fat mass (C), lean mass (D), 5-hours fasting blood glucose level (E, left), glucose tolerance during the intraperitoneal glucose tolerance test (ipGTT; F, left) and corresponding area under the curve (F, right), and blood glucose levels during the intraperitoneal insulin tolerance test (ipITT; G) in control (CTL) and fructose-fed (FRU) adult male mice. (H, I) Memory performance. Exploration time of the moved object in the object location memory test (H) and latency to reach the hidden platform during acquisition in the Morris water maze (MWM) (I, left). Time spent in the different zones of the Morris water maze during the probe trial (I, right), including quadrants adjacent to the original platform location and the target quadrant. Data are presented as mean ± SEM; * *p < 0.05,* ** ***p < 0.01,*** *** p < 0.001, unpaired t-test vs. CTL; ## p < 0.01, ### p < 0.001, diet effect in two-way ANOVA; $ p < 0.05, $$ p < 0.01 *vs.* chance (50% in the OLM; 25%in the MWM); ns, non-significant, p > 0.05.

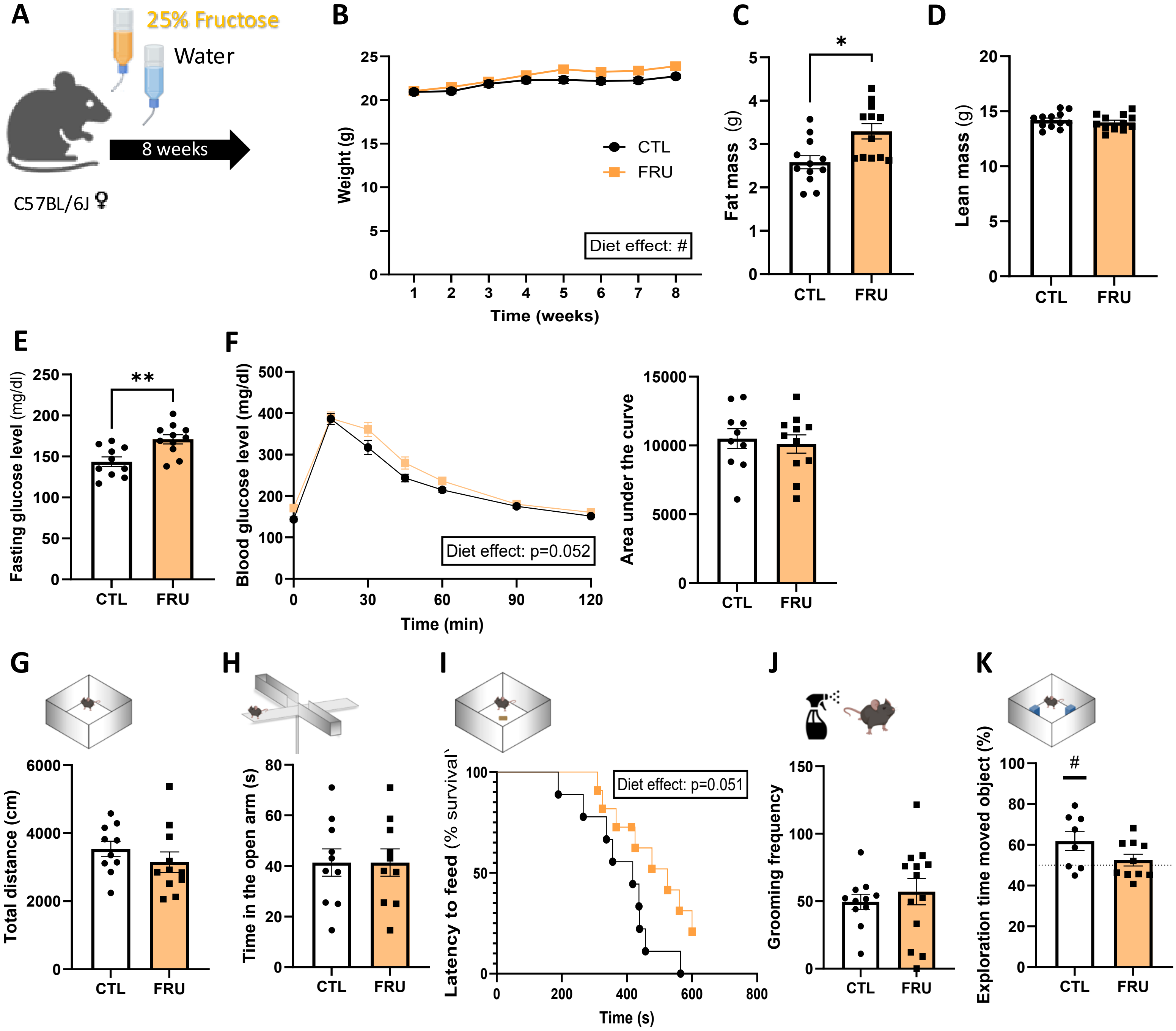


<u>**Supp. Figure 2:**</u> **Metabolic and behavioral effects of chronic fructose intake in adult female mice.** (A) Experimental design: adult C57BL/6J female mice were exposed to 25% fructose in drinking water or control water for 8 weeks. (B–F) Metabolic parameters. Body weight gain (B), fat mass (C), lean mass (D), 5-hours fasting blood glucose level (E), glucose tolerance during the ipGTT (F, left panel) and corresponding area under the curve (F, right panel) in control (CTL) and fructose-fed (FRU) mice. (G–K) Emotional cognitive behaviors. Total locomotion (locomotor activity) in the openfield test (G), time spent in open arms of the elevated plus maze (H), latency to feed (% survival) in the novelty-suppressed feeding test (I), grooming frequency in the splash test (J), and exploration time in the object location memory test (K). Data are presented as mean ± SEM; *p < 0.05, **p < 0.01; unpaired t-test *vs.* CTL; # p < 0.05; one-sample t-test *vs.* 50%

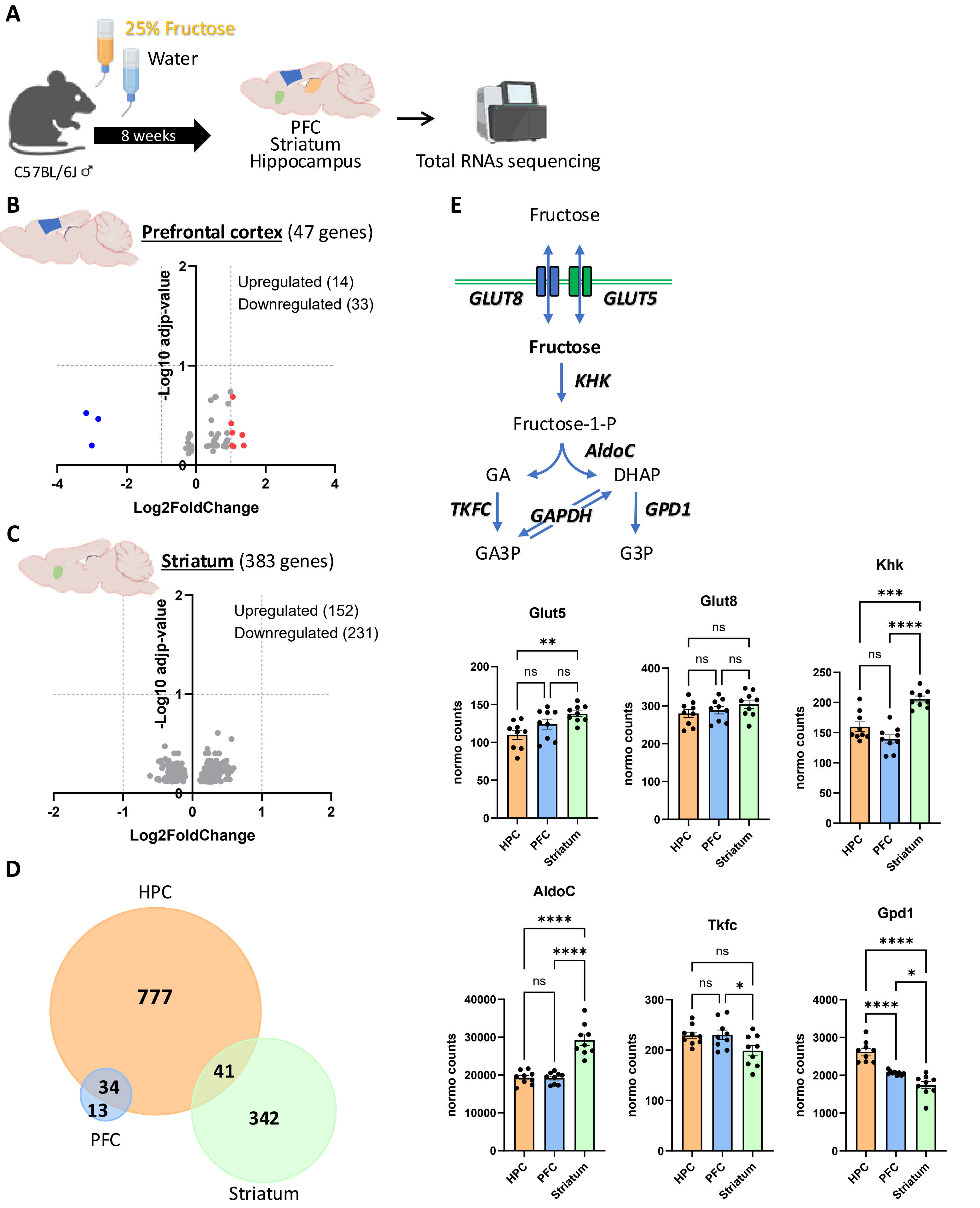

A
25% Fructose
Water
8 weeks
C57BL/6J ♂
PFC
Striatum
Hippocampus
Total RNAs sequencing
B
Prefrontal cortex (47 genes)
Upregulated (14)
Downregulated (33)
-Log10 adjp-value
Log2FoldChange
C
Striatum (383 genes)
Upregulated (152)
Downregulated (231)
-Log10 adjp-value
Log2FoldChange
D
HPC
777
34
13
41
342
PFC
Striatum
E
Fructose
GLUT8
GLUT5
Fructose
KHK
Fructose-1-P
AldoC
GA
DHAP
TKFC
GAPDH
GPD1
GA3P
G3P
Glut5
Glut8
Khk
AldoC
Tkfc
Gpd1
normo counts
HPC
PFC
Striatum
ns

**<u>Supp. Figure 3:</u> Fructose-induced transcriptional changes and regional expression of genes of fructose transport and metabolism genes.** A. Experimental design: RNAseq analysis performed in the hippocampus (HPC), prefrontal cortex (PFC) and striatum of adult male mice exposed to 25% fructose in drinking water or control water for 8 weeks. (B, C) Volcano plots showing differentially expressed genes (DEGs) in the PFC (B) and striatum (C). Upregulated genes above 1 Log2FoldChange (Log2FC) are highlighted in red and downregulated genes below -1 Log2FC in blue, with the total number of DEGs indicated for each brain region. (D) Venn diagram illustrating the overlap of DEGs between HPC, PFC, and striatum, highlighting the large number of hippocampal-specific changes and the limited overlap across regions. (E) Schematic representation of fructose transport and metabolism genes, including GLUT5- and GLUT8-mediated fructose uptake and key enzymes of the fructose metabolism (Khk, AldoC, Tkfc, Gpd1), together with RNA-seq–derived expression levels of the corresponding genes in the HPC, PFC, and striatum. Bar graphs display normalized counts (mean ± SEM). * $p < 0.05$, ** $p < 0.01$, *** $p < 0.001$ **** $p < 0.0001$; One-way ANOVA followed by Tukey's multiple comparisons test.

**A**

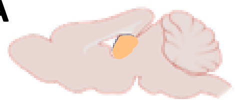

Hippocampus

| | Cell type | DEGs overlapping | Odds ratio | p-value |
|---|---|---|---|---|
| **All DEGs** | Astrocytes | 42 / 852 | 1.08 | 0.334 |
| | Neurons | 37 / 852 | 0.74 | 0.968 |
| | Microglia | 27 / 852 | 0.64 | 0.993 |
| | Oligodendrocytes | 23 / 852 | 0.48 | 1.000 |
| | OPCs | 4 / 852 | 0.22 | 1.000 |
| | Endothelial | 15 / 852 | 0.33 | 1.000 |
| **Up-regulated DEGs** | Neurons | 35 / 543 | 1.15 | 0.236 |
| | Astrocytes | 25 / 543 | 1.00 | 0.522 |
| | OPCs | 4 / 543 | 0.36 | 0.995 |
| | Oligodendrocytes | 11 / 543 | 0.36 | 1.000 |
| | Endothelial | 9 / 543 | 0.32 | 1.000 |
| | Microglia | 5 / 543 | 0.18 | 1.000 |
| **Down-regulated DEGs** | Microglia | 22 / 309 | 1.55 | **0.040** |
| | Astrocytes | 17 / 309 | 1.22 | 0.253 |
| | Oligodendrocytes | 12 / 309 | 0.72 | 0.895 |
| | Endothelial | 6 / 309 | 0.38 | 0.998 |
| | Neurons | 2 / 309 | 0.11 | 1.000 |
| | OPCs | 0 / 309 | 0.00 | 1.000 |

**B**

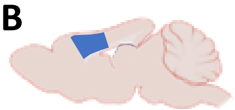

Prefrontal cortex

| | Cell type | DEGs overlapping | Odds ratio | p-value |
|---|---|---|---|---|
| **All DEGs** | Neurons | 7/47 | 3.07 | **1,27E-02** |
| | Astrocytes | 4/47 | 2.02 | 1,52E-01 |
| | Endothelial | 1/47 | 0.43 | 9,04E-01 |
| | Oligodendrocytes | 1/47 | 0.41 | 9,13E-01 |
| | Microglia | 0/47 | 0 | 1.000 |
| | OPCs | 0/47 | 0 | 1.000 |
| **Up-regulated DEGs** | Endothelial | 1/14 | 1.51 | 5,01E-01 |
| | Oligodendrocytes | 1/14 | 1.45 | 5,16E-01 |
| | Neurons | 1/14 | 1.34 | 5,42E-01 |
| | Astrocytes | 0/14 | 0 | 1.000 |
| | Microglia | 0/14 | 0 | 1.000 |
| | OPCs | 0/14 | 0 | 1.000 |
| **Down-regulated DEGs** | Neurons | 6/33 | 3.89 | **7,88E-03** |
| | Astrocytes | 4/33 | 3.0 | 5,56E-02 |
| | Endothelial | 0/33 | 0 | 1.000 |
| | Microglia | 0/33 | 0 | 1.000 |
| | OPCs | 0/33 | 0 | 1.000 |
| | Oligodendrocytes | 0/33 | 0 | 1.000 |

**C**

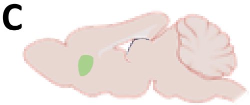

Striatum

| | Cell type | DEGs overlapping | Odds ratio | p-value |
|---|---|---|---|---|
| **All DEGs** | Astrocytes | 39/383 | 2.49 | **1,90E-06** |
| | Neurons | 40/383 | 1.8 | **9,18E-04** |
| | Endothelial/Vascular | 13/383 | 0.83 | 7,74E-01 |
| | OPCs | 3/383 | 0.67 | 8,19E-01 |
| | Oligodendrocytes | 18/383 | 0.76 | 8,99E-01 |
| | Microglia | 6/383 | 0.43 | 9,94E-01 |
| **Up-regulated DEGs** | Astrocytes | 37/152 | 7.2 | **1,56E-17** |
| | Endothelial/Vascular | 11/152 | 1.88 | **4,26E-02** |
| | Oligodendrocytes | 15/152 | 1.71 | **4,30E-02** |
| | OPCs | 2/152 | 1.15 | 5,22E-01 |
| | Microglia | 5/152 | 0.93 | 6,27E-01 |
| | Neurons | 0/152 | 0 | 1.000 |
| **Down-regulated DEGs** | Neurons | 40/231 | 3.28 | **3,05E-09** |
| | OPCs | 1/231 | 0.37 | 9,32E-01 |
| | Endothelial/Vascular | 2/231 | 0.21 | 9,99E-01 |
| | Astrocytes | 2/231 | 0.18 | 1.000 |
| | Microglia | 1/231 | 0.12 | 1.000 |
| | Oligodendrocytes | 3/231 | 0.2 | 1.000 |

**Supp. Figure 4: Cell type enrichment analysis of fructose-induced transcriptional changes across brain regions.** RNA-seq was performed on hippocampus (A), prefrontal cortex (B), and striatum (C) from adult male mice exposed to control or 25% fructose for 8 weeks. Differentially expressed genes (DEGs) were identified and overlapped with published cell type–specific gene signatures to assess enrichment in astrocytes, neurons, endothelial/vascular cells, oligodendrocytes, oligodendrocyte precursor cells (OPCs), and microglia. Tables show, for each region, the number of overlapping DEGs relative to the total DEGs, together with odds ratios and p-values for all DEGs, upregulated DEGs, and downregulated DEGs. $p < 0.05$ are indicated in **bold**.

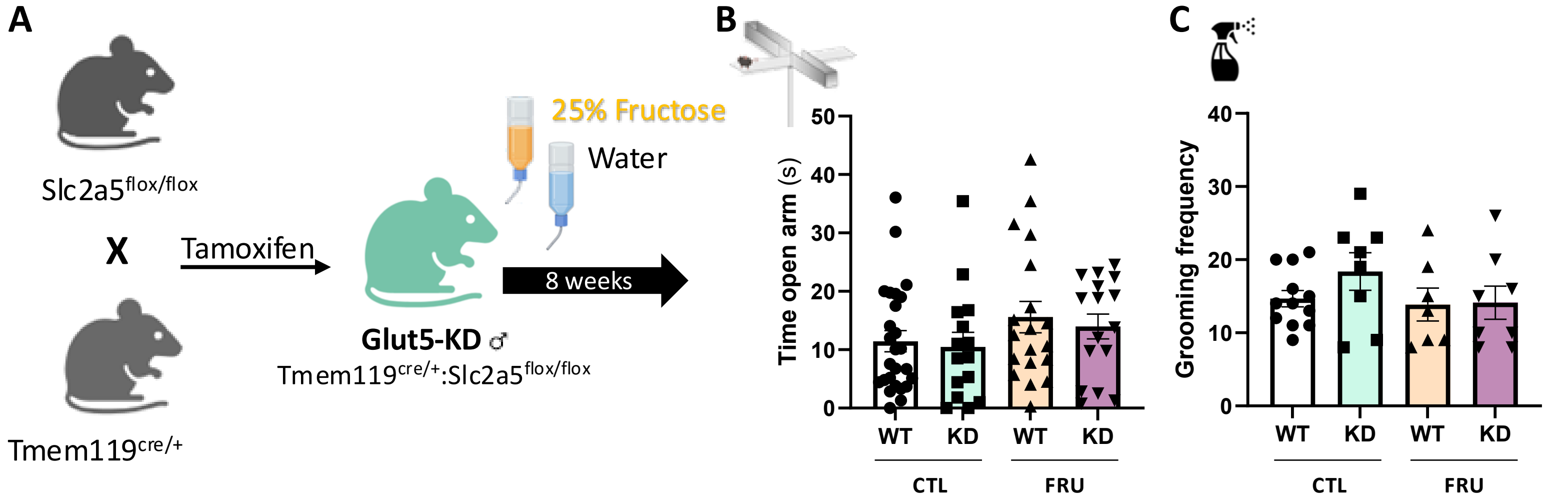


**Supp. Figure 5: Emotional behaviors in microglial Glut5-KD mice after chronic fructose exposure.** (A) Experimental design: Slc2a5$^{flox/flox}$ mice were crossed with Tmem119$^{creERT2}$ mice to generate microglia-specific Glut5 knockdown (Glut5-KD; Tmem119$^{cre/+}$:Slc2a5$^{flox/flox}$) and wild-type (WT) littermates (Slc2a5$^{flox/flox}$ cre negative). Adult male WT and Glut5-KD mice were treated with tamoxifen to induce Cre-mediated Glut5 deletion in microglia, then exposed to either control drinking water (CTL) or 25% fructose solution (FRU) for 8 weeks. (B–C) Emotional behaviors. Time spent in open arms of the elevated plus maze (B) and grooming frequency in the splash test (C) in WT and Glut5-KD mice. Data are presented as mean ± SEM; statistical analysis was performed using two-way ANOVA.

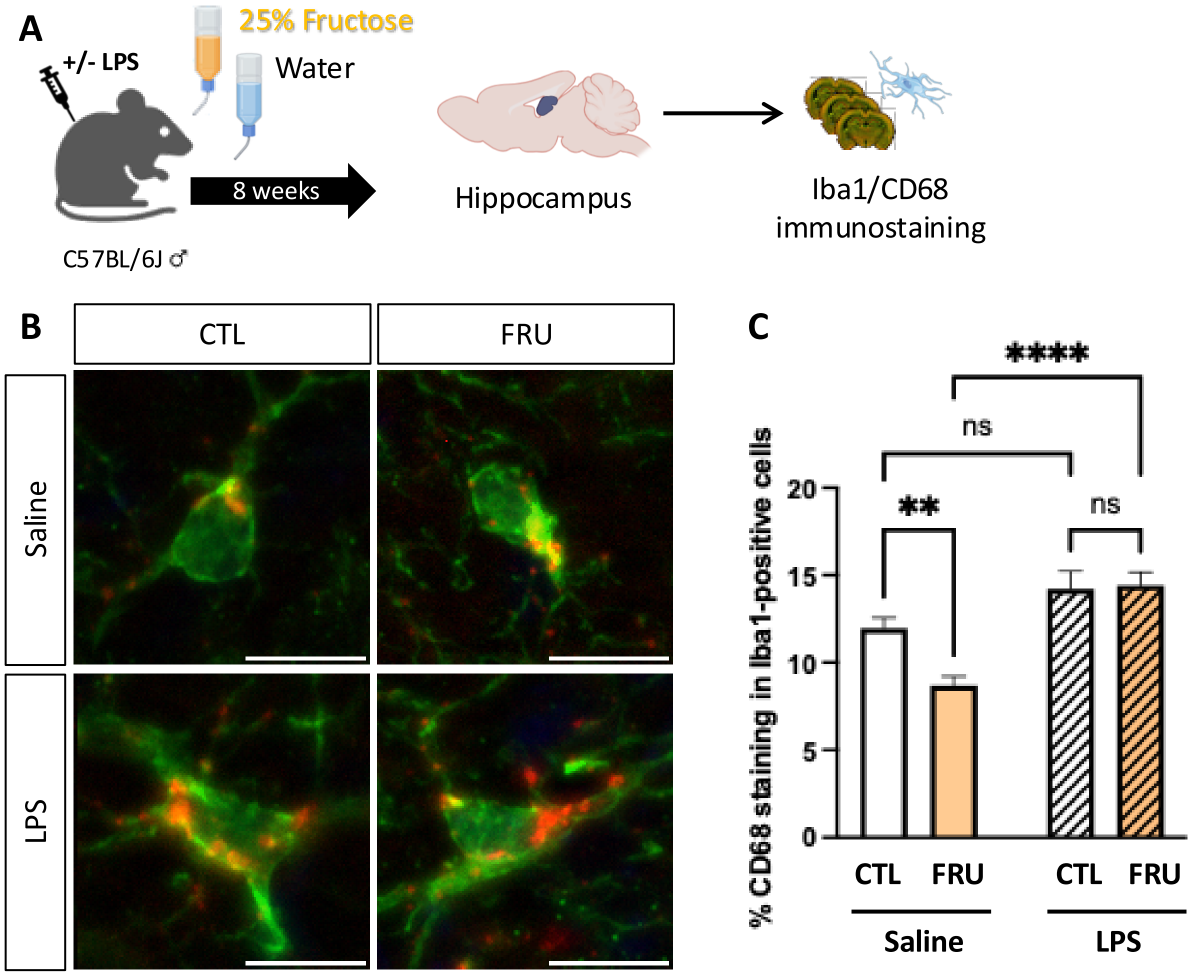


<u>**Supp. Figure 6:**</u> **CD68 immunoreactivity in fructose fed mice treated or not with LPS.** (A) Experimental design: adult C57BL/6J male mice were exposed to 25% fructose in drinking water or control water (CTL) for 8 weeks before being treated with LPS (250 µg/kg). (B, C) Representative images (B, Scale bar 10 µm) and quantification (C) of the % of CD68 immunostaining area in Iba1-positive cells in the hippocampus measured 24-hours post-LPS treatment. Data are presented as mean ± SEM; ns $p > 0.05$. ** $p < 0.01$; **** $p < 0.0001$ (two-way ANOVA followed by Tukey post-hoc test).

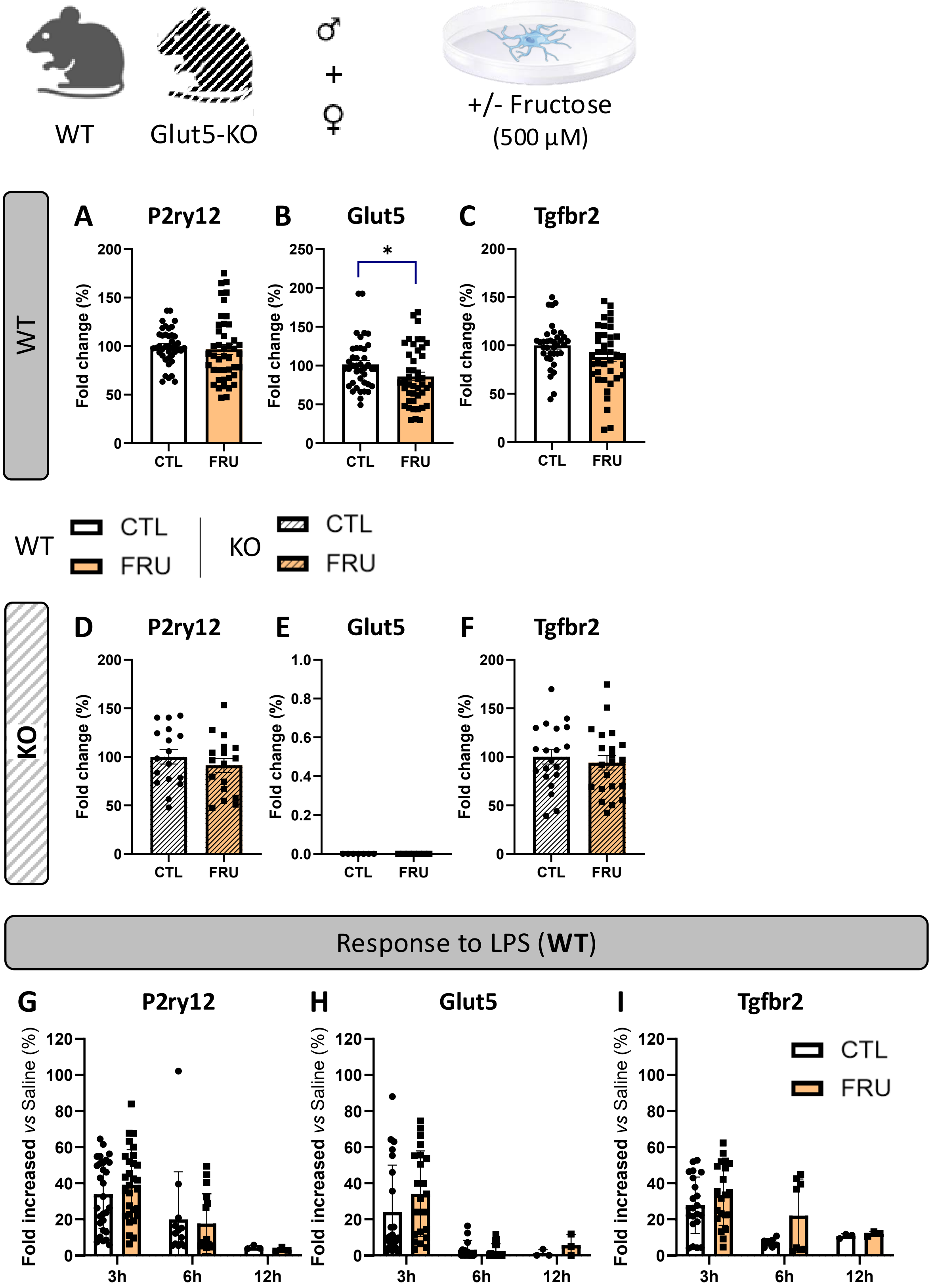


**Supp. Figure 7: Effects of fructose on microglial homeostatic gene expression *in vitro*.** Primary microglia were isolated from adult wild-type (WT) or whole-body Glut5 knockout (Glut5-KO) male and female mice and cultured with or without fructose (500 µM) for 5 days before being treated with LPS (100 ng/mL). (A–F) RT-qPCR analysis of homeostatic microglial markers in microglia from WT (A-C) or Glut5-KO (D-F) mice treated with vehicle (CTL) or fructose (FRU): P2ry12 (A, D), Glut5 (B, E), and Tgfbr2 (C, F), expressed as fold change (%) relative to CTL. (G–I) Response of microglia cultured from WT mice to LPS. Time course of P2ry12 (G), Glut5 (H) and Tgfbr2 (I) expression following LPS stimulation, represented as percentage fold change vs. unstimulated saline treated cells at 3, 6, and 12 h, in the presence (FRU) or absence (CTL) of fructose. Data are presented as mean ± SEM; *p < 0.05 *vs.* CTL (unpaired t-test).
